# CASM potentiates STING-driven NFκB signaling in immune cells

**DOI:** 10.64898/2026.01.21.700774

**Authors:** Isis Benoit-Lizon, Cassandre Pignol, Kirsty Hooper, Caroline Truntzer, Abdelmnim Radoua, Ludivine Dal Zuffo, Julie Lucas, Quentin Frenger, Emy Ponsardin, Clarisse Cornebise, Isabelle Cantaloube, Ameetha Ratier, Romain Aucagne, François Hermetet, Sayyed Hamed Shahoei, Erik R. Nelson, Emeric Limagne, Christian Poüs, Claudine Deloménie, Pauline Soulas-Sprauel, Frédéric Gros, Frédérique Végran, Carmen Garrido, François Ghiringhelli, Catherine Paul, Oliver Florey, Elise Jacquin, Lionel Apetoh

**Author notes:** Contributed equally (Elise Jacquin, Lionel Apetoh). INSERM U1151, CNRS UMR8253, Institut Necker Enfants Malades, Université Paris Cité, Paris, France. The Jackson Laboratory for Genomic Medicine, Farmington, CT, USA.

## Abstract

Stimulator of Interferon Gene (STING), a key player of antimicrobial immune responses, has emerged as a promising target to mitigate inflammation and cancer. Following STING activation, proinflammatory molecules and type I Interferons (IFN) are released thus favoring the establishment of effective immune responses and adaptive immunity. Autophagy has been proposed to negatively regulate STING signaling. While STING activation drives microtubule-associated proteins 1A/1B light chain 3B (hereafter referred to as LC3) lipidation, the underlying mechanisms and functional consequences remain however incompletely defined. Especially, the consequences of STING-associated Conjugation of autophagy related (ATG) 8 to Single Membranes (CASM) in the control of immune responses remain elusive. Using innate and adaptive cells specifically inactivated for autophagy or CASM, we found that STING agonists primarily trigger CASM over autophagy. While STING-associated autophagy exerts negative feedback on the STING pathway and downstream type I IFN and pro-inflammatory responses, with different underlying molecular mechanisms between immune cells, STING-driven CASM potentiates NFκB-associated TNF production. These results overall uncover a new function of CASM and underscore the relevance of both CASM and autophagy in shaping STING signaling

## Introduction

STING (Stimulator of interferon genes protein), is an endoplasmic reticulum membrane-resident protein involved in cytoplasmic DNA sensing (Ishikawa & Barber, 2008). Initially related to antiviral immune response, the cGAS (Cyclic GMP-AMP synthase)-STING pathway has been extensively studied in the context of inflammation and cancer (Decout et al., 2021; Zhu et al., 2019). Mechanistically, STING downstream signaling involves the IRF3 (Interferon regulatory factor 3) and RELA/p65 (nuclear factor kappa B (NF-κB) transcription factor p65) which drive the production of type I Interferons (IFN), pro-inflammatory cytokines such as IL-6 (Interleukin-6) and TNF (Tumor necrosis factor) as well as chemo-attracting molecules such as CCL5 (C-C motif chemokine 5) and CXCL10 (C-X-C motif chemokine 10) (Zhang et al., 2025; Zhu et al., 2019). While STING triggering in myeloid cells favors their activation, STING engagement in lymphocytes also has functional consequences. For instance, we demonstrated that intrinsic STING activation can directly modulate CD4^+^ T cell effector and antitumor functions (Benoit-Lizon et al., 2022). Although STING is involved in broad range of immune related process and pathologies (Chen & Xu, 2023) and despite the development of various new STING targeting molecules (Haag et al., 2018; Jacoby et al., 2025), a deeper understanding of the STING pathway regulation at both cellular and molecular levels is needed for the design of STING-based therapies.

The lipidation of the ATG (Autophagy-related) 8 protein family member MAP1LC3/LC3 (microtubule associated protein1 light chain 3, hereafter referred to as LC3) has rapidly emerged as a common event following STING activation (Gui et al., 2019; Konno et al., 2013; Liu et al., 2019; Prabakaran et al., 2018). Together with other ATG8 proteins GABARAP (Gamma-aminobutyric acid receptor-associated protein) and GABARAPL1 and L2 (GABARAP like 1 and like 2), LC3 proteins are key components of the autophagy machinery. These ubiquitin-like proteins are conjugated to the phosphatidylethanolamines of forming autophagosomes during the catabolic process of macro-autophagy (hereafter referred to as autophagy) where they contribute to autophagic cargo selection and double membrane autophagosome formation. LC3 lipidation has thus been widely used as a marker of autophagy. Importantly, alternative lipidation of ATG8 proteins to single membrane compartments instead of double-membrane autophagosomes has been evidenced (Florey et al., 2011; Sanjuan et al., 2007). Distinct from conventional autophagy, this non-canonical autophagy process, recently renamed Conjugation of Atg8 proteins to Single Membrane (CASM), occurs upon various macroendocytic processes such as entosis (Florey et al., 2011), phagocytosis of apoptotic cell or pathogens (LC3-associated phagocytosis, LAP) and endocytosis (LC3-associated endocytosis, LANDO) (Heckmann et al., 2020). CASM is also triggered by pharmacological agents, viroporins or ion channel agonists targeting acidic intracellular vesicles as well as endolysosome damage (Cross et al., 2023; Durgan & Florey, 2022; Fletcher et al., 2018; Florey et al., 2015; Jacquin et al., 2017; Ulferts et al., 2021). Mechanistically, CASM requires the LC3 lipidation machinery (ATG3-ATG5-ATG7-ATG10-ATG12-ATG16L1) but is independent of the upstream autophagy regulator mTOR and the autophagy initiation complex (Florey et al., 2015; Florey et al., 2011; Jacquin et al., 2017). The vacuolar H^+^-ATPases proton pump (V-ATPase) is a key regulator of CASM which recruits the core LC3 lipidation protein ATG16L1 to single membranes by direct interaction with its C-terminal WD40 domain (Hooper et al., 2022; Timimi et al., 2022). Deletion of ATG16L1 WD40 domain or a single point mutation of its K490 residue is sufficient to render cells incompetent for CASM, while they remain competent for autophagosome formation, allowing to distinguish between the two pathways (Fletcher et al., 2018).

LC3 lipidation following cGAS or STING activation has previously been associated with autophagosome formation which in turn facilitates the clearance of cytosolic DNA or mediates the degradation of STING itself or the downstream components of the STING pathway TBK1 (TANK binding kinase 1, Serine/threonine-protein kinase) and IRF3 (Jiang et al., 2018; Liu et al., 2019; Pan et al., 2023; Prabakaran et al., 2018; Wu et al., 2021; Xie et al., 2022). STING-driven LC3 lipidation can also occur on single-membrane perinuclear vesicles (Saitoh et al., 2009) and independently of ULK1 (unc-51 like autophagy activating kinase 1), RB1CC1/FIP200 (RB1 inducible coiled-coil 1), ATG13, PIK3C3/VPS34 (phosphatidylinositol 3-kinase catalytic subunit type 3), Beclin-1, WIPI2 (WD repeat domain phosphoinositide-interacting protein 2) and autophagy receptors (Fischer et al., 2020; Gui et al., 2019; Liu et al., 2019). Using cancer cells and murine embryonic fibroblasts (MEFs) deficient for autophagy components Fischer et al. described for the first time STING-associated CASM, that they named V-ATPase–ATG16L1—induced LC3B lipidation (VAIL) (Fischer et al., 2020). Identification of STING’s proton channel function has clarified the molecular mechanisms linking STING activation and LC3 conjugation to single-membrane acidic vesicles (B. Liu et al., 2023; Xun et al., 2024). Meanwhile, the exploration of STING-associated CASM functional consequences has evidenced its role in lysosomal homeostasis maintenance (Bentley-DeSousa et al., 2025; Eguchi et al., 2024; Huang et al., 2025; Lv et al., 2024; Tang et al., 2025; Xu et al., 2025). Despites these recent advances, a deeper understanding of how STING-driven LC3 lipidation can modulate immune functions is still needed.

In this study, we explore the molecular mechanisms underlying STING-driven LC3 lipidation in immune cells, which are essential effectors of STING signaling during infection but also inflammatory diseases and cancers (Knox et al., 2025). We also use autophagy or CASM-incompetent innate and adaptive immune cells to discriminate between the two pathways and show that STING agonists not only trigger autophagy but also CASM *in vitro* and *in vivo*. Finally, we find that while STING-driven autophagy exerts negative feedback on type I IFN and pro-inflammatory responses, STING-driven CASM rather supports NFκB signaling. Our results therefore clarify the interplay between STING and autophagy proteins in immune cells and unravel a novel immune-related function for CASM.

## Results

### STING activation mainly drives CASM in innate and adaptive immune cells

STING agonists have been shown to stimulate LC3 lipidation *in vitro* in various cell lines (Fischer et al., 2020; Gui et al., 2019; Hooper et al., 2022; Liu et al., 2019; Prabakaran et al., 2018; Saitoh et al., 2009). We tested the consequences of STING activation on LC3 lipidation in immune cells *in vivo* in two different pathological contexts. First, we used a STING-associated vasculopathy with onset in infancy (SAVI) mouse model with a heterozygous STING gain-of-function mutation V154M/WT (STING^V154M/WT^) driving its constitutive activation (Bouis et al., 2019). We observed an increase in LC3 lipidation in splenocytes isolated from mice harboring the V154M/WT mutation compared to WT control littermates. This demonstrated that constitutive STING activation could drive LC3 lipidation in immune cells *in vivo*, in the absence of any exogenous stimulation (**Figure 1A**). Given that STING can also be activated in tumors, we tested whether the mouse STING agonist DMXAA (DMX) could stimulate LC3 lipidation in tumor-infiltrating lymphocytes (TILS). To refine the origin of LC3 lipidation driven by STING activation, we took advantage of a mouse model of *Rb1cc1*/*Fip200* conditional deletion in CD4^+^ T cells (*Rb1cc1*^flox/flox^:Cd4^Cre^ referred to as *ΔFip200*), in which CD4^+^ T cells are incompetent for autophagy as illustrated by the absence of Bafilomycin A1 (BafA1) effect on LC3 lipidation and their accumulation of the autophagy receptor p62 (**Figure S1A**). MC38 tumor-bearing *ΔFip200* mice were treated with intratumoral (*i.t.*) injection of STING ligand (DMX) or control (DMSO) before isolation of CD4^+^ TILS and flow cytometry-based LC3 lipidation measurement. Treatment with DMX increased LC3 lipidation in CD4^+^ TILS (**Figure 1B**) indicating that STING agonists used for cancer therapy could trigger LC3 lipidation in tumor infiltrating lymphocytes *in vivo*. The observation that this lipidation occurred independently of FIP200 suggested that STING activation stimulated CASM in lymphocytes of the TME.

**Figure 1.**
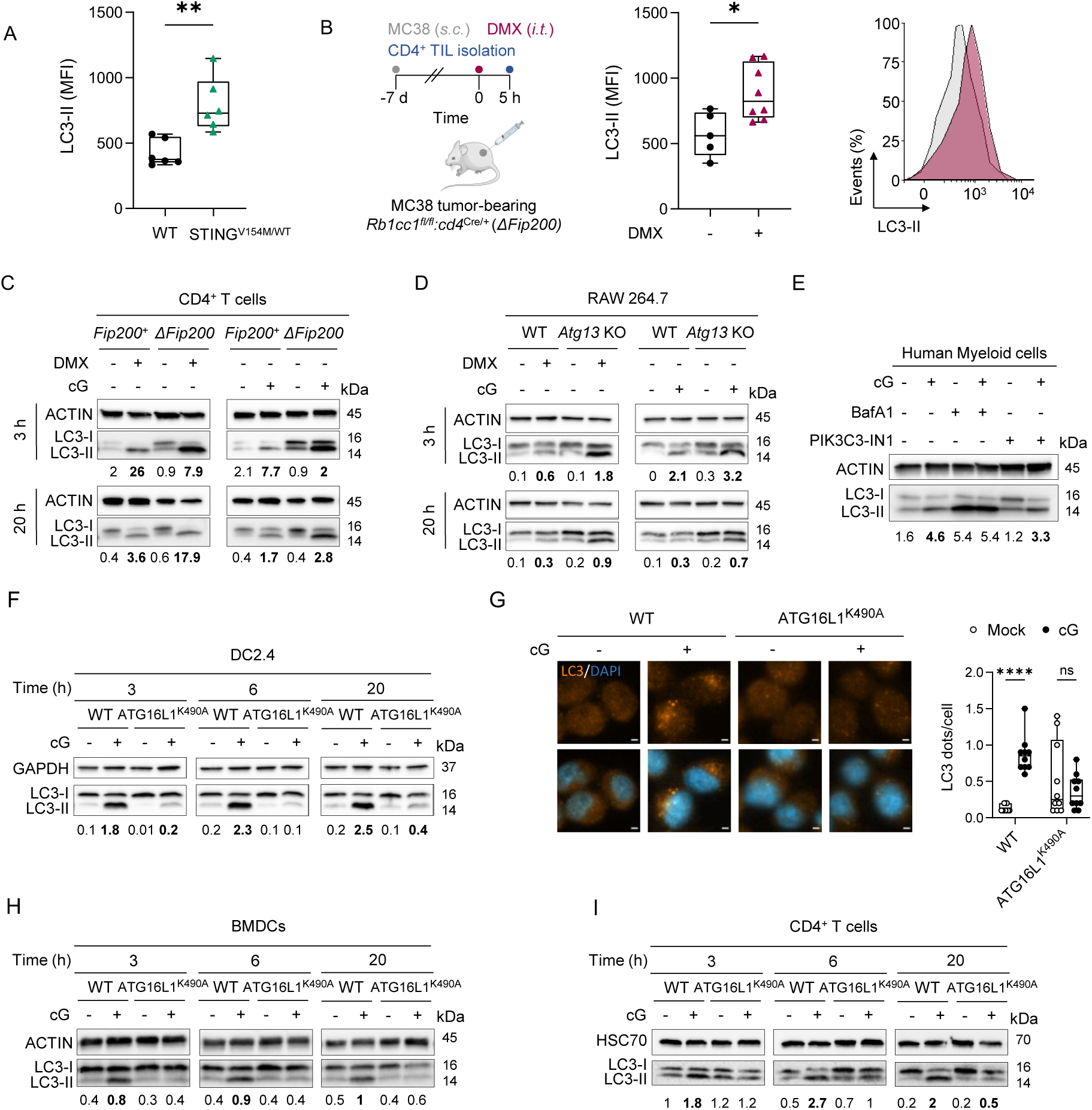
STING activation mainly drives autophagosome-independent LC3-lipidation that requires ATG16L1 lysine 490. LC3 lipidation (LC3-II FITC MFI) analyzed by flow cytometry (**A and B**) in splenocytes from WT or STING^V154M/WT^ mutated mice (**A**) or tumor infiltrating CD4^+^ T cells from *Fip200^fl/fl^:cd4^Cre/+^* mice, treated or not with i.t. injection of DMX (250 µg) for 5 h (**B**), n= 5-8 mice from 3 independent experiments, P values (*p<0.05, **p<0.01) determined by unpaired t-test. LC3 lipidation analyzed by western blot (**C-F**) in *Fip200*^+^ or *ΔFip200* CD4^+^ T cells (**C**) stimulated or not *ex vivo* with DMXAA or 2’3’-cGAMP (DMX or cG) for 3 or 20 h, representative of n=2-3 independent experiments; in WT or *Atg13* KO RAW264.7 macrophages (**D**) stimulated or not with DMX or cG for 3 or 20 h, representative of n=2-3 independent experiments; in adherent/myeloid cells isolated from human PBMCs (**E**) stimulated or not with cG for 3 h with or without PI3KC3/VPS34 inhibitor (PI3KC3-IN1) or Bafilomycin A1, representative of n=3 donors; in WT or ATG16L1^K490A^ DC2.4 cells (**F**), stimulated or not with cG for 3, 6 or 20 h, representative of n=3 independent experiments. LC3 lipidation analyzed by immunofluorescence in WT or ATG16L1^K490A^ DC2.4 cells (**G**) stimulated or not with cG for 3 h, quantification of LC3 dots from ten fields of view, each containing >200 cells, **** P < 0.0001, and representative images, scale bar, 2 µm. LC3 lipidation analyzed by western blot (**H and I**) in WT or ATG16L1^K490A^ BMDCs (**H**) or CD4^+^ T cells (**I**), stimulated or not with cG for 3, 6 or 20 h, representative of 2 to 6 independent experiments. LC3-II:LC3-I ratios are indicated below western blot images and increases in LC3-II:LC3-I ratios, compared to control conditions, are highlighted in bold.

We then sought to dissect the mechanisms driving LC3 lipidation in innate and adaptive immune cells. After demonstrating that STING-driven LC3-lipidation requires the components of the LC3 lipidation machinery ATG5 and ATG16L1 (**Figure S1B to D**) we tested the requirement of the autophagy pre-initiation complex proteins FIP200 and ATG13 which are essential for autophagosome formation but dispensable for CASM (Jacquin et al., 2017) (**Figure S1A and E**). LC3 lipidation was observed in *ΔFip200* CD4^+^ T cells stimulated *ex vivo* **(Figure 1C**) and in *Atg13* KO RAW264.7 macrophages stimulated *in vitro* (**Figure 1D**) with the STING agonists DMX and 2’-3’-cGAMP (cG). These STING agonists thus triggered autophagosome-independent LC3 lipidation upon short term and prolonged stimulation in both lymphoid and myeloid cells. We also used a drug-based approach in which Bafilomycin A1 (BafA1) increases LC3 lipidation associated with canonical autophagy, but impairs CASM due to its inhibitory action on V-ATPase; whereas the PIK3C3/VPS34 inhibitor 1 (PIK3C3-IN1) inhibits autophagosome formation, but does not impair most forms of CASM (Jacquin et al., 2017). This approach was validated in RAW264.7 murine macrophages (**Figure S1F**) and used in myeloid cells isolated from healthy donor peripheral blood mononuclear cells (PBMCs). In these autophagy-competent cells stimulated *ex vivo*, combination of cG with BafA1 did not result in a further increase in LC3 lipidation compared to BafA1 alone, while PIK3C3 inhibition did not abrogate cG-driven LC3 lipidation (**Figure 1E**), suggesting again that STING activation rapidly triggered CASM not only in mouse but also in human primary immune cells.

To definitively show that STING activation drove CASM in immune cells, we used a unique genetic system to distinguish between autophagy and CASM. A single point mutation of the K490 amino-acid in ATG16L1 C-terminal WD40 domain (referred to as ATG16L1^K490A^) inactivates CASM while cells remain capable of autophagosome formation. We thus generated CASM-incompetent mouse dendritic cells (DC2.4) using a CRISPR knock-in approach and combined pharmacological autophagy and CASM modulators to functionally validate our model. Both the V-ATPase inhibitor BafA1 and the ionophore Monensin (Mon) blocks autophagic flux by raising lysosomal pH but only Mon also stimulates CASM. We thus combined these two drugs with PIK3C3-IN1 which prevents autophagosome formation and only allows for CASM-associated LC3 lipidation (**Figure S1G**). Contrary to ATG16L1 WT cells, DC2.4 cells expressing ATG16L1^K490A^ showed reduced LC3 lipidation upon stimulation with Mon combined with PIK3C3-IN1 compared to Mon alone indicating their inability to conjugate LC3 to single-membrane vesicles (Fletcher et al., 2018; Hooper et al., 2022). They remain however able to lipidate LC3 to autophagosomes as illustrated by the PIK3C3/VPS34-dependent increase in LC3-II:LC3-I ratio upon BafA1 treatment in both WT and ATG16L1^K490A^ cells (**Figure S1G**). Western-blot and immunofluorescence analyses showed that LC3 lipidation triggered by STING activation with cG was strongly reduced in CASM-incompetent DC2.4 cells compared to WT cells (**Figure 1F and G**), supporting STING-driven CASM. Similar results were obtained in *Atg16l1* KO RAW264.7 macrophages stably re-expressing 7WT ATG16L1 or ATG16L1^K490A^ (**Figure S1H**). Likewise, while STING activation with cG stimulated LC3 lipidation in primary myeloid (BMDCs) and lymphoid (CD4^+^ T cells) isolated from WT mice, it failed to induce or induced only mild LC3 lipidation in primary immune cells isolated from CASM-incompetent (AG16L1^K490A^) littermates (**Figure 1H and I**). Altogether, these data strongly supported STING-associated CASM in myeloid and lymphoid immune cells.

### STING-driven CASM requires STING but not downstream components of the STING pathway or Rubicon

We then tested the requirement of STING pathway components for STING ligand - driven CASM. Given that short term STING activation mainly drives CASM in primary mouse immune cells (**Figure. 1H and I**, 3 h timepoint), we differentiated BMDCs and isolated CD4^+^ T cells from STING-deficient (*Sting* KO) or WT mice and stimulated them *ex vivo* with STING agonists for 3 h. We used different types of STING agonists including the flavone-acetic acid-based drug DMX and the cyclic dinucleotides cG and cdi-AMP or cdi-GMP respectively produced by mammalian cells and bacteria. While the mouse STING agonist DMX, the mammalian cyclic dinucleotide cG and the bacterial-derived cdi-AMP and cdi-GMP all triggered CASM in WT BMDCs (**Figure 2A**) and CD4^+^ T cells (**Figure 2B**), they all failed to drive LC3 lipidation in the corresponding STING-deficient immune cells. Similar results were obtained following prolonged stimulation with STING agonists (**Figure S2A and B**). STING expression by innate and adaptive immune cells was thus required for CASM induction by these agonists. Overall, different types of STING agonists activated CASM in immune cells.

**Figure 2.**
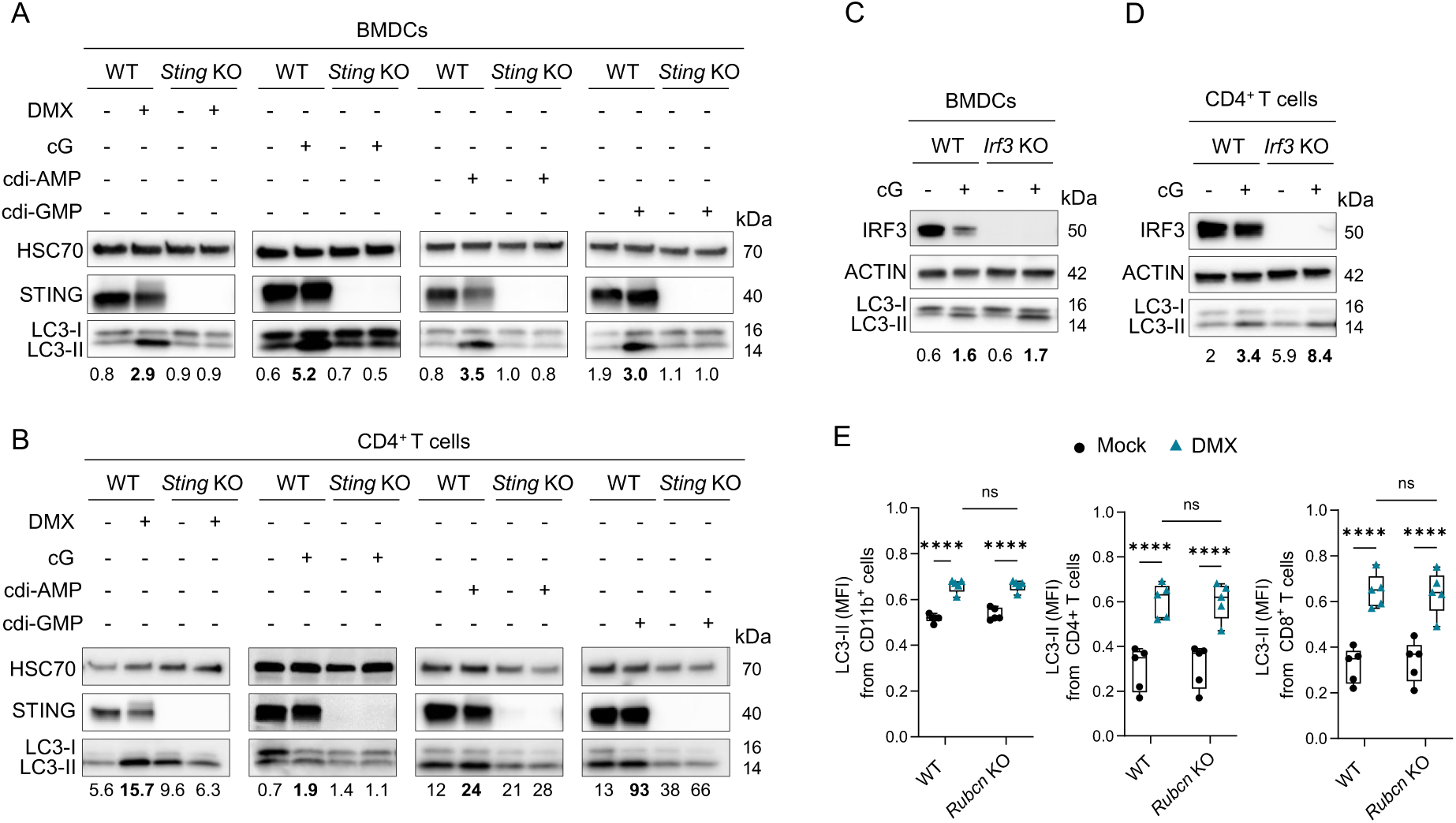
STING-driven CASM is induced by various STING ligands and requires STING but not IRF3 or RUBICON. LC3 lipidation analyzed by western blot (**A to D)** in WT or *Sting* KO BMDCs (**A**) or CD4^+^ T cells (**B**), stimulated or not with DMX, cG, cdi-AMP or cdi-GMP for 3 h, representative of n=2 to 5 independent experiments; in WT or *Irf3* KO BMDCs (**C**) or CD4^+^ T cells (**D**), stimulated or not with cG for 3 h, representative of n=3 independent experiments. LC3 lipidation (LC3-FITC MFI, normalized to Bafilomycin A1 condition) analyzed by flow cytometry (**E**) in WT or *Rubcn* KO CD11b^+^ cells, CD4^+^ or CD8^+^ T cells, stimulated or not with DMX for 3 h, n= 5 mice from 3 independent experiments, P values (*p<0.05, **p<0.01) determined by two-way ANOVA. LC3-II:LC3-I ratios are indicated below western blot images and increases in LC3-II:LC3-I ratios, compared to control conditions, are highlighted in bold.

We then tested the requirement of the STING pathway downstream components TBK1 and IRF3 for STING-associated CASM. We thus respectively combined STING agonists with an inhibitor of the kinase TBK1 (BX795) in RAW 264.7 macrophages (**Figure S2C**) and stimulated BMDCs and CD4^+^ T cells isolated from IRF3-deficient mice (*Irf3* KO) with cG (**Figure 2C and D, Figure S2D and E**). While TBK1 inhibition with BX795 impaired IRF3 phosphorylation induced by cG and DMX, it did not suppress LC3 lipidation (**Figure S2C**). Furthermore, STING activation with cG triggered LC3 lipidation in IRF3-deficient cells (**Figure 2C and D; Figure S2D and E**). STING-driven CASM thus required neither TBK1 nor IRF3 as previously suggested for STING-associated LC3 lipidation in non-immune cell lines (Gui et al., 2019; Liu et al., 2019).

Rubicon (RUN domain and cysteine-rich domain containing, Beclin 1–interacting protein) acts within a PIK3C3/VPS34 complex responsible for phosphatidylinositol-3-phosphate (PtdIns3P) formation during phagocytosis and for the inhibition of autophagy (Magné & Green, 2022). Deletion of Rubicon has been shown to significantly inhibit CASM during LC3-associated phagocytosis and LC3-associated endocytosis (Cunha et al., 2018; Heckmann et al., 2019). We thus tested if Rubicon was required for STING-associated CASM by stimulating primary immune cells from WT and Rubicon-deficient (*Rubcn* KO) mice with the STING agonist DMX before measuring LC3 lipidation by flow cytometry. We found that DMX stimulated LC3 lipidation to the same extent in WT and *Rubcn* KO myeloid (Cd11b^+^) and lymphoid (CD4^+^ and CD8^+^) cells indicating that STING-associated CASM occurring in immune cells is independent of Rubicon (**Figure 2E**).

### STING-driven autophagy negatively regulates STING signaling

We observed that LC3 lipidation triggered by the STING agonist cG is not always fully abrogated in CASM-incompetent cells, suggesting that STING activation can also stimulate autophagosome formation. Indeed, residual CASM-independent LC3 lipidation was observed in CASM-incompetent DC2.4 (**Figure 1F and G**), RAW264.7 macrophages (**Figure S1H**) as early as 3 h after STING activation and in CASM-incompetent primary CD4^+^ T cells after prolonged STING activation with cG (**Figure 1I**), illustrating the activation of autophagy. We thus sought to assess the specific consequences of STING-associated autophagy on immune cell response to STING activation by using autophagy-incompetent myeloid and lymphoid immune cells lacking ATG13 or FIP200 respectively, two components of the autophagy pre-initiation complex essential for autophagosome formation.

We first compared the expression changes of STING-associated genes between autophagy-competent and -incompetent immune cells following stimulation with cG. We found that STING activation further stimulated the expression of type I IFN and pro-inflammatory cytokine encoding genes as well as IFN-stimulated gene (ISG) expression in autophagy-incompetent immune cells, compared to control cells. Stimulation of *Atg13* KO RAW 264.7 macrophages with cG led to a significantly higher increase in *Ifnb1*, *Tnf*, *Ifit1*, *Ifit2*, *Mx2*, *Cxcl10* and *Ccl5* mRNA levels (**Figure 3A; Figure S3A and B**) as well as IFN-β and CXCL10 secretion (**Figure 3B**) compared to WT cells. Similarly, stimulation of *ΔFip200* CD4^+^ T cells with cG led to a significantly higher increase in *Ifnb1, Il6* and *Ccl5* mRNA levels and a light elevation in *Tnf*, *Ifit1*, *Ifit2*, *Mx2* and *Cxcl10* mRNA levels (**Figure 3C, S3C and D**) compared to autophagy competent *Fip200*^+^ CD4^+^ T cells. We accordingly observed a further increase in IFN-β, CXCL10 and IL6 secretion (**Figure 3D**). Autophagy thus negatively regulated both myeloid and lymphoid immune cell responses to STING activation.

**Figure 3.**
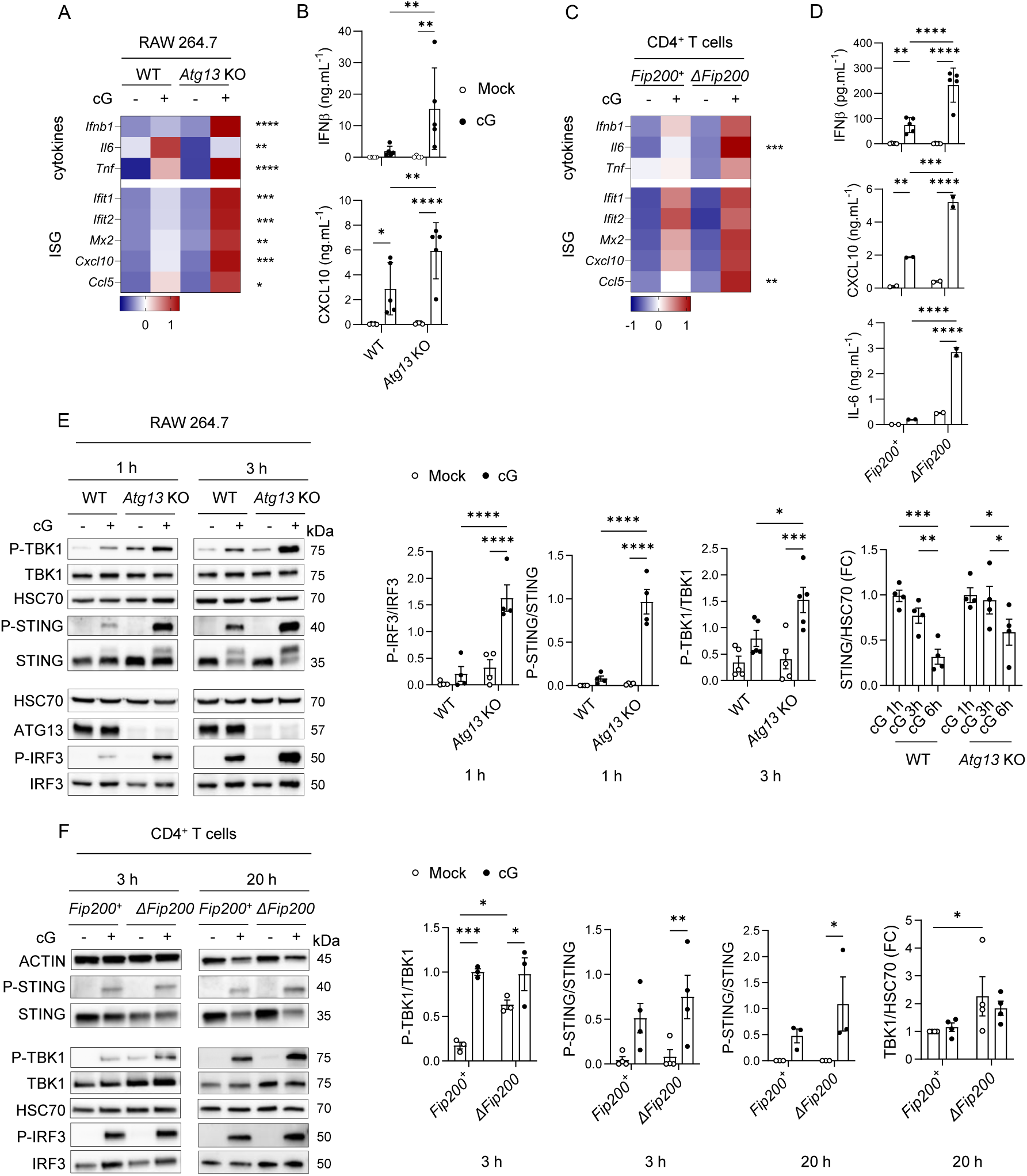
STING-associated immune responses are upregulated in autophagy-incompetent macrophages and T cells. (**A**) Heatmap representing indicated cytokine and ISG mRNA expression (Z-score) determined by RTqPCR from WT and *Atg13* KO RAW264.7 macrophages stimulated or not with cG for 3 h. (**B**) IFN-β and CXCL10 secretion measured by ELISA from WT and *Atg13* KO RAW264.7 macrophages stimulated or not with cG for 3 h or 6 h respectively. (**C**) Heatmap representing indicated cytokine and ISG mRNA expression (Z-score) determined by RTqPCR from *Fip200^+^* and *ΔFip200* CD4^+^ T cells stimulated or not with cG for 20 h. (**A and C**) Pooled data from 4 independent experiments, P values (*p<0.05, **p<0.01, ***p<0.001, ****p<0.0001, ns not represented) determined by two-way ANOVA from each target relative expression. (**D**) IFN-β, CXCL10 and IL-6 secretion measured by ELISA from *Fip200*^+^ and *ΔFip200* CD4^+^ T cells stimulated or not with cG 20 h. (**B and D**), pooled data (Mean +/- SD) from 2 to 5 independent experiments, P values (*p<0.05, **p<0.01, ***p<0.001, ****p<0.0001, ns not represented) determined by two-way ANOVA. (**E and F**) Western blot analyses of indicated proteins and corresponding relevant quantifications from WT and *Atg13* KO RAW264.7 macrophages (**E**) or *Fip200*^+^ and *ΔFip200* CD4^+^ T cells (**F**) stimulated or not with cG for indicated time, pooled data (Mean +/- SEM) from 3 to 5 independent experiments, P values (*p<0.05, **p<0.01, ***p<0.001, ****p<0.0001, ns not represented) determined by two-way ANOVA.

We investigated the underlying molecular mechanisms by assessing the phosphorylation and levels of STING pathway components. Increased phosphorylation of STING, TBK1 and IRF3 was observed in *Atg13* KO RAW 264.7 macrophages upon stimulation with cG compared to WT cells. Our data also suggested that activation-induced STING degradation was impaired in *Atg13* KO RAW 264.7 macrophages (**Figure 3E, S3E**). In myeloid cells, STING-associated autophagosome formation thus contributed to STING degradation following activation, exerting negative feedback on the pathway as suggested by previous studies (Pan et al., 2023; Prabakaran et al., 2018). In *ΔFip200* CD4^+^ T cells, STING and IRF3 phosphorylation tended to be higher than in *Fip200*^+^ CD4^+^ T cells following STING activation (**Figure 3F; Figure S3F**). This further increase in STING signaling went along with a higher level of TBK1 phosphorylation at baseline and TBK1 accumulation over time in autophagy-deficient cells associated with the phosphorylation and activation of a higher amount of STING and IRF3 proteins thus sustaining STING signaling (**Figure 3F; Figure S3F**). Furthermore, prolonged STING activation in autophagy-incompetent CD4^+^ T cells rather seemed associated with a reduced activation-induced degradation of IRF3, than an impairment of activation-induced degradation of STING. Overall, these data indicated that STING-associated autophagy negatively regulated STING signaling in immune cells with difference in kinetics and molecular mechanisms between myeloid and lymphoid cell types.

### STING-driven CASM is required for robust RELA/p65 activation and subsequent TNF production in innate immune cells

To explore the functional consequences of STING-associated CASM in immune cells, we first compared the transcriptional changes induced by cG in BMDCs differentiated from WT and CASM-incompetent ATG16L1^K490A^ mice. Gene set enrichment analysis (GSEA) revealed that pathways associated with innate immune response, positive regulation of cytokine production and chemokine receptor binding were significantly enriched in WT BMDCs but not CASM-incompetent BMDCs (**Figure 4A; Figure S4A**). This suggested that CASM could support robust immune response following STING activation. To test this hypothesis, we measured STING-associated gene expression in WT and ATG16L1^K490A^ BMDCs stimulated or not with cG. Type I IFN-encoding genes and ISG were induced following STING activation to the same extent in WT and ATG16L1^K490A^ BMDCs (**Figure 4B, Figure S4B**) similarly to IFN-β secretion (**Figure 4C**). STING signaling also appeared similarly activated in WT and ATG16L1^K490A^ BMDCs as illustrated by the comparable levels of IRF3 phosphorylation and activation-induced STING degradation (**Figure 4D**). Similar analyses conducted in WT and ATG16L1^K490A^ CD4^+^ T cells suggested a stronger or faster response of CASM-incompetent cells compared to control cells illustrated by a further increase in STING-associated mRNA (**Figure S4C**), but this was neither associated with a further increase in IFN-β secretion nor with further activation of STING signaling (**Figure S4D and E**).

**Figure 4:**
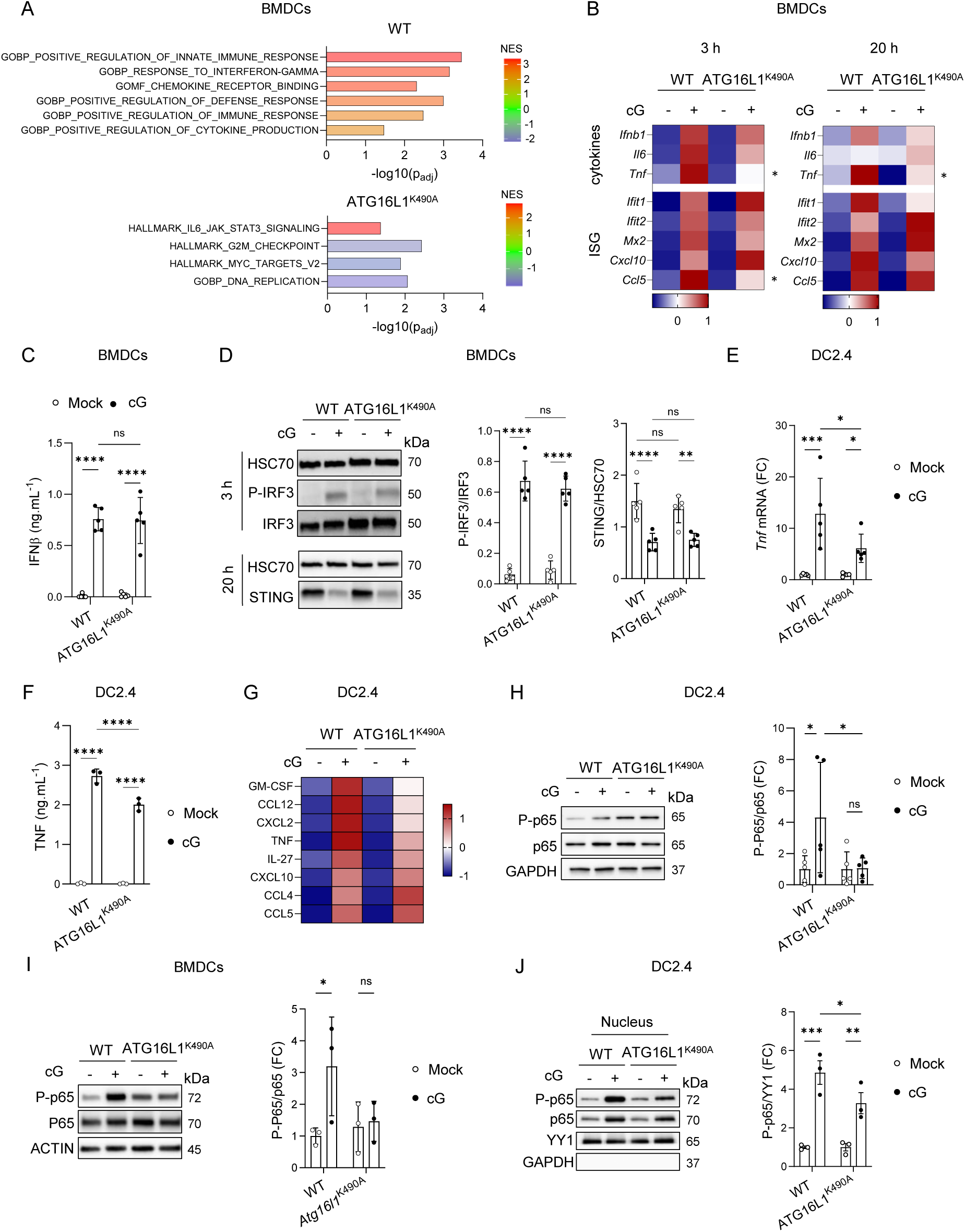
STING-driven CASM regulates RELA/p65-induced inflammatory response in myeloid cells. (**A**) GSEA (Gene set enrichment analysis; Normalized Enrichment Score (NES) & -log10 (p_adj_)) obtained from RNA sequencing analysis of WT or ATG16L1^K490A^ BMDCs stimulated or not with cG for 3 h. (**B**) Heatmap representing indicated cytokine and ISG mRNA expression (Z-score) determined by RTqPCR from WT or ATG16L1^K490A^ BMDCs stimulated or not with cG for 3 or 20 h, pooled data (Mean) from n=6-7 mice from 3 independent experiments, P values (*p<0.05) determined by two-way ANOVA from each target relative expression. IFN-β secretion measured by ELISA (**C**) or western blot analysis of indicated proteins and corresponding quantifications (**D**) from WT or ATG16L1^K490A^ BMDCs stimulated or not with cG for 3 (**D**) or 20 h (**C and D**), pooled data (Mean +/- SD) from n=5 mice from 2 independent experiments, P values (**p<0.01, ***p<0.001, ****p<0.0001) determined by two-way ANOVA. *Tnfa* mRNA expression determined by RTqPCR (Fold Change FC; Normalized to mean of each control condition) (**E**) and TNF-α secretion measured by ELISA (**F**) from WT or ATG16L1^K490A^ DC2.4 cells stimulated or not with cG for 6 h, pooled data (Mean +/- SD) from 3 to 5 independent experiments, P values (*p<0.05, ***p<0.001, ****p<0.0001) determined by two-way ANOVA. (**G**) Heatmap representing indicated molecule secretion (Z-score) measured by Cytokine Array from WT or ATG16L1^K490A^ DC2.4 cells stimulated or not with cG for 6 h, pooled data from 2 independent experiments. Western blot analysis of indicated proteins from total (**H and I**) or nuclear (**J**) extracts and corresponding quantifications (Fold Change FC, normalized to mean of WT mouse control condition) from WT or ATG16L1^K490A^ DC2.4 cells (**H and J**) or BMDCs (**I**) stimulated or not with cG for 3 h, pooled data (Mean & SD) from 5 independent experiments (**H**) or from n=3 mice (**I and J**), P values (*p<0.05, **p<0.01) determined by two-way ANOVA.

Contrary to type I IFN, *Tnf* mRNA induction was reduced in ATG16L1^K490A^ BMDCs compared to WT cells upon stimulation with cG (**Figure 4B; Figure S4F**). This dampened TNF response was also observed in CASM-incompetent DC2.4 cells at both mRNA and secretion levels (**Figure 4E and F**). Using cytokine array, we found that not only TNF but also GM-CSF, CCL12 and CXCL2 were less secreted following STING activation from CASM-incompetent DC2.4 cells than WT cells (**Figure 4G; Figure S4G**). We also observed reduced *Cxcl2* and *Ccl12* mRNA (**Figure S4H**). Expression of these cytokines and chemokines being transcriptionally regulated by NF-κB (Burke et al., 2014; Lopes et al., 2018), we compared NF-κB signaling between WT and ATG16L1^K490A^ DC2.4 cells upon STING activation with cG. STING-associated phosphorylation of RELA/p65 (Ser 536) was reduced in CASM-incompetent DC2.4 cells compared to WT (**Figure 4H**) which was also observed in primary BMDCs (**Figure 4I**). Nuclear localization of phosphorylated RELA/p65 (Ser 536) upon stimulation of DC2.4 cells with cG was also lower in CASM-incompetent cells compared to WT control cells (**Figure 4J**). Overall, these data showed that CASM did not regulate IRF3 signaling and type I IFN production in immune cells but rather sustains the NF-κB response downstream STING activation, thereby potentiating RELA/p65 phosphorylation and subsequent TNF production.

## Discussion

While an interplay between the autophagy machinery and the STING was described shortly after STING identification (Saitoh et al., 2009), whether STING activation drives conventional autophagy or CASM and how autophagy protein activity can in turn modulate cell response to STING activation has remained unclear and context-dependent. Here, we found that STING activation drives both autophagy and CASM in immune cells, leading to distinct outcomes. The first association between STING and autophagy proteins already reported unexpected findings. Indeed, Saitoh et al. revealed that upon activation of the cGAS-STING pathway with dsDNA, STING translocated to punctate structures characterized as single-membrane vesicles rather than double membrane autophagosomes (Saitoh et al., 2009). Follow-up studies accordingly evidenced LC3 lipidation following STING activation (Fischer et al., 2020; Gui et al., 2019; Liu et al., 2019; Prabakaran et al., 2018). However, the underlying molecular mechanisms were not clearly defined and often characterized as a non-conventional form of autophagy, independent of the upstream autophagy components mTOR, ULK1/2, ATG13, FIP200, Beclin1, VPS34 and ATG9A but accompanied by STING targeting to autophagosomes through its interaction with the autophagy receptor p62 or directly with LC3 via its LIR motif (Gui et al., 2019; Liu et al., 2019). The single-membrane nature of the intracellular compartments where STING translocates following activation as well as the absence of requirement of the autophagy initiation and PIK3C3 complexes were reminiscent of CASM (Durgan et al., 2021; Florey et al., 2011). Using genetically engineered cancer cells and fibroblasts, Fischer et al. described V-ATPase–ATG16L1—induced LC3B lipidation (VAIL) which occurred on single-membrane vesicles, independently of FIP200, WIPIs and autophagy receptors. VAIL was insensitive to VPS34 inhibition whereas it required ATG16L1 WD40 domain and V-ATPase and was inhibited by the Salmonella typhimurium virulence, SopF (Fischer et al., 2020). Despite this alternative name, STING-associated VAIL is thus CASM mediated by V-ATPase subunit association and ATG16L1 recruitment as shown by Hooper et al. in ATG16L1 KO mouse macrophages re-expressing WT or ATG16L1K490A (Hooper et al., 2022). Combining different mouse and cell models of autophagy or CASM inactivation, our approach allowed to discriminate between the two pathways and showed that CASM appears as the primary source of LC3 lipidation following STING activation in innate and adaptive immune cells (**Figure 1F to I**), occurring a few hours after STING agonist addition and maintained over time. Indeed, STING-driven LC3 lipidation was scarcely altered by RB1CC1/FIP200 and ATG13 loss whereas it was strongly inhibited in CASM-incompetent cells expressing ATG16L1^K490A^ protein. CASM associated with LC3-associated phagocytosis and LC3-associated endocytosis was shown to require PtdIns3P and ROS production as well as Rubicon activity (Cunha et al., 2018; Heckmann et al., 2019) contrary to CASM activated by ionophores, lysosomotropic agents (Jacquin et al., 2017) or TRPLM1 agonists (Goodwin et al., 2021). Our data showing similar LC3 lipidation following STING activation in Rubicon-deficient primary cells and their WT counterparts indicate that STING-driven CASM occurs independently of Rubicon and reinforces the idea that Rubicon is not required for all form of CASM (Durgan & Florey, 2022). Importantly, we show for the first time that FIP200-independent CASM is activated in tumor-infiltrating lymphocytes following STING agonist administration to tumor-bearing mice. Thus, the increased LC3 lipidation observed in mice carrying the gain-of-function V154M/WT STING mutation could at least partly be associated with CASM. This evidence of STING-driven CASM *in vivo*, together with our observation of VPS34 inhibition-insensitive STING-driven CASM in human myeloid primary cells isolated from donor blood, raise an essential question regarding its functional consequences while STING agonists are under clinical evaluation for cancer treatment (Jacoby et al., 2025; Richter et al., 2023; Samson & Ablasser, 2022)

Besides CASM, STING activation was also suggested to drive autophagy in the seminal work of Saitoh et al. who observed STING colocalization with the autophagy receptor p62 (Saitoh et al., 2009). This was confirmed by Prabakaran et al. who observed increased cytoplasmic levels of double-membrane autophagosomes, ULK1 serine 555 phosphorylation as well as LC3 lipidation following STING activation in MEFs. Additional experiments conducted in the human monocytic cancer cell line THP-1 led the authors to a conclusion of p62-dependent selective autophagy activation (Prabakaran et al., 2018). Here, we show that STING activation in immune cells can stimulate autophagy in parallel with CASM, providing a second source of LC3 lipidation, with differences between cell types (**Figure 1F to I, S1H**). Autophagy induction upon STING activation was previously proposed to exert negative feedback on the STING pathway (Ji et al., 2023; Liu et al., 2019; Prabakaran et al., 2018). We thus investigated the functional consequences of STING-driven autophagy in immune cells and the differences in underlying mechanisms between innate and adaptive immune cells. Autophagy suppression by *Atg13* or *Rb1cc1/Fip200* deletion leads to increased type I IFN and pro-inflammatory responses to STING activation in both innate and adaptive immune cells. However, our investigation of STING signaling revealed different underlying molecular mechanisms. In RAW264.7 macrophages, STING signaling, illustrated by STING, TBK1 and IRF3 phosphorylation levels following STING activation, is markedly increased and stimulation-induced STING degradation is reduced in autophagy-incompetent (*Atg13* KO cells). In these cells, autophagy thus exerts negative feedback on the STING pathway through autophagy-mediated degradation of STING, as previously reported in cancer cells (THP-1, HeLa) and in MEFs (Liu et al., 2019; Pan et al., 2023; Prabakaran et al., 2018). Stimulation-induced STING degradation was not completely abrogated in autophagy-incompetent cells which supports others studies describing lysosomal (Chu et al., 2021; Gonugunta et al., 2017; Konno et al., 2013; Zhu et al., 2022), ERAD-mediated (Ji et al., 2023), ESCRT-dependent (Balka et al., 2023; Gentili et al., 2023) or proteasome-mediated (Gonugunta et al., 2017; Liu et al., 2019; Prabakaran et al., 2018) STING degradation and confirms that autophagy is not the only process regulating STING activity. Alternatively, the increased STING signaling observed in autophagy-incompetent (*ΔFip200*) T cells appears associated with a basal increased in TBK1 phosphorylation, TBK1 accumulation over time, and to a minor extent, a decrease in IRF3 degradation following STING activation. These data fits with studies showing how selective autophagy can mediate TBK1 and IRF3 degradation and dampen downstream type I IFN responses (Jiang et al., 2018; Wu et al., 2021; Xie et al., 2022; Zhao et al., 2022). Collectively, our work and previous studies demonstrate that STING signaling is regulated by multiple processes, including the autophagic degradation of STING or downstream components of the STING pathway, but also of cGAS (Chen et al., 2016; Liang et al., 2014; Zhao et al., 2021), and cytosolic DNA (Gui et al., 2019; Liang et al., 2014; Y. Liu et al., 2023), with differences between cell types, even among immune cells. These observations should be considered for the development of therapies targeting STING-driven inflammation and immune responses. For instance, inhibitors of lysosomal activity and/or autophagy enhance STING agonist antitumor activity *in vivo* in mouse melanoma. Gonugunta et al. found that combining the V-ATPase inhibitor Bafilomycin A1 with a low dose of the STING agonist cG allows for tumor growth inhibition (Gonugunta et al., 2017). Similarly, Yu et al. combined an inhibitor of PIK3C3/VPS34 named SB02024, which blocks autophagosome formation and affects late endosomes and lysosomes, with the cG analog ADU-S100. They found that PIK3C3/VPS34 inhibition improved ADU-S100 antitumor efficacy and mouse survival (Yu et al., 2024). Taken together, these data suggest that targeting autophagy could be a interesting way to improve the efficacy of STING agonists for cancer treatment while clinical trials have evidenced limitations (Hines et al., 2023; Lu et al., 2023; Schmid et al., 2024).

Since its discovery, non-canonical autophagy or CASM has received growing attention leading to the identification of many new stimuli including ion channels such as TRPML1 or STING (Durgan & Florey, 2022). Indeed, the recently identified proton channel function of STING shed light on the molecular mechanisms driving CASM upon STING activation. STING conformational change upon agonist binding was proposed to enable proton leakage from Golgi-derived acidic vesicles (Huang et al., 2025; B. Liu et al., 2023; Xun et al., 2024). In response, V-ATPase subunit association in these deacidified vesicle membranes recruits ATG16L1 leading to CASM (Fischer et al., 2020; Hooper et al., 2022; Huang et al., 2025). The functional consequences of STING-driven CASM remain less understood. Recent studies have identified a role for STING-driven CASM in endosomal and lysosomal homeostasis through different pathways. First, the lipidation of GABARAPs following STING activation drives the TFEB/TFE3-dependent transcription of lysosome and autophagy-related genes (Huang et al., 2025; Lv et al., 2024; Xu et al., 2025) similar to what has been observed following activation of the lysosomal ion channel TRPML1 (Goodwin et al., 2021). Second, CASM-associated GABARAP lipidation maintains lysosome homeostasis in an LRKK2/Rab10-dependent manner (Bentley-DeSousa et al., 2025; Eguchi et al., 2024). Again, this appears as a common feature of CASM, induced by various CASM-inducers creating endolysosome perturbation. Finally, GABARAP lipidation upon endolysosome perturbation, induced by the STING agonists MSA2 and diABZI prevents the accumulation of enlarged and probably damaged vacuoles through ESCRT recruitment (Huang et al., 2025). Although this role for CASM in endolysosomal maintenance and homeostasis seems to be specific to GABARAP lipidation, it suggests that STING-driven CASM can create new signaling hubs similar to what has been previously suggested following TLR activation (Acharya et al., 2016; Hayashi et al., 2018). However, no clear role for CASM in STING-related signaling and immune functions has been identified. Previous work has shown that CASM is dispensable for STING-associated cell death (Xun et al., 2024), NLRP3 activation (B. Liu et al., 2023) and type I Interferon production (Gui et al., 2019; Tang et al., 2025; Xun et al., 2024). Here, we made similar observation regarding the absence of a role for CASM in STING-dependent regulation of IRF3 signaling and type I IFN production. We found however that STING-driven CASM may be important for NF-κB-mediated inflammation. Indeed, CASM-incompetent dendritic cells have reduced RELA/p65 activation following STING activation with cG (**Figure 4H and I**), leading to a reduced production of TNF and the pro-inflammatory chemokines CXCL2 and CCL12 (**Figure 4E to G, S4G and H**). We thus propose that STING-driven CASM can amplify NF-κB signaling and the production of downstream immune effectors. In their recent study, Fischer et al. found that STING activation stimulates HOIP-mediated M1-ubiquitin (M1-Ub) chain formation which then activates NF-κB-related gene expression. While they showed that CASM inactivation with SopF disturbs the localization of HOIP and ubiquitin, it does not impair the formation of M1-Ub chains. Furthermore, inhibition of STING-driven LC3 lipidation by ATG16L1 deletion impaired neither type I IFN nor NF-κB-related gene expression. Conversely, a mild, but significant, increase in *TNF* and *TNFAIP3* mRNA levels was observed in ATG16L1 KO THP-1 cells compared to WT control cells after prolonged exposure to the STING agonist diABZI (Fischer et al., 2020). ATG16L1 deletion inactivating both CASM and autophagy-associated LC3 lipidation, this increase in NF-κB-related gene expression could reflect the loss of autophagy-mediated negative feedback on the STING pathway rather than a function of CASM. While downstream activation of IRF3 and type I IFN signaling upon STING activation has been extensively studied, STING-associated NF-κB signaling still deserves some mechanistic investigations. Our data, showing that STING-driven CASM fine-tunes NF-κB signaling and TNF production but not IRF3 signaling and type I IFNs, may seem unexpected since TBK1 has been initially found to phosphorylate both IRF3 and RELA/p65 following STING activation (Abe & Barber, 2014). However, recent reports have shown that TBK1 and its homolog IKKɛ can act redundantly downstream STING activation (Balka et al., 2020) and that IKKɛ activity is largely restricted by its expression level. Indeed, while TBK1 is ubiquitously expressed, IKKɛ expression is limited to some immune cell populations (Venkatraman et al., 2024). Understanding how NF-κB signaling is regulated following STING activation has remained challenging, especially in immune cells in which its activation is frequent, even in the absence of danger signal. Our findings suggest that STING-driven CASM could serve as a signaling platform to recruit and activate kinase complexes, for efficient phosphorylation of p65, and uncover new crosstalk between STING and NF-κB signaling.

The consequences of CASM on inflammation and immune response have rarely been explored *in vivo*. Wang et al. found that CASM limits protects mice from lethal respiratory inflammation following Influenza A virus (IAV) infection (Wang et al., 2021). These observations were supported by a recent study showing that STING-induced CASM can facilitate pathogen clearance (Xu et al., 2025). In a tumor context, CASM occurring upon phagocytosis (LAP) has been shown to limit STING activation and downstream type I IFN response, possibly by preventing the release of DNA from engulfed apoptotic cells in the cytosol. This results in the polarization of tumor-associated macrophage (TAM) polarization towards a immunosuppressive and protumor phenotype (Cunha et al., 2018). Depending on the context, CASM could thus regulate STING-associated immune responses not only by preventing STING activation but also by supporting an efficient production of NF-κB-associated pro-inflammatory molecules.

Overall, our work shows that STING activation triggers both CASM and autophagy in immune cells with different functional consequences. While STING-associated autophagy exerts negative feedback on the STING pathway through different underlying mechanisms, STING-driven CASM rather supports NF-κB signaling. Identifying a new role for CASM in immune response, this study underscores the significance of this pathway in STING-associated inflammation and immunity.

## Materials and Methods

### Mice

Wild type (WT) female C57BL/6 mice were purchased from Charles River laboratories (France). *Sting1*^-/-^ mice (*Sting* KO; provided by Pr. Bernhard Ryffel), *Irf3*^-/-^ (*Irf3* KO; RBRC00858 - RIKEN), CD4-Cre mice were purchased from JAX (stock numbers 17336), *Atg5*^flox/flox^ mice were purchased from RIKEN (RBRC02975), *Rb1cc1*^flox/flox^ were provided by Dr Jun-Lin Guan and Geoffrey Pinski (University of Cincinnati). Mice were all bred at the TAAM (Typage et archivage d’animaux modèles, UAR44 – CNRS, Orléans, France). CD4-Cre mice were crossed with *Atg5*^flox/flox^ or mice *Rb1cc1*^flox/flox^ to obtain mice conditionally deficient for ATG5 or FIP200 (respectively Δ*Atg5* and Δ*Fip200*) and their control littermates. STING V154M/WT (STING^V154M/WT^) and Rubicon^-/-^ (*Rubcn* KO) mice were bred and provided by Pr Soulas-Sprauel and Dr Gros (UMR 1109 INSERM/Université de Strasbourg, Strasbourg, France). ATG16L1^K490A^ mice were bred and mouse organs were provided by Dr Florey (The Babraham Institute, Cambridge, UK).

All animals were bred and maintained according to both the FELASA and the Animal Experimental Ethics Committee Guidelines (University of Burgundy and TAAM, Orléans, France). The Ethics Committee for Animal Welfare of the University of Burgundy and the French Ministry of Higher Education and Research approved all animal experiments (references APAFIS #32752-2021082512009877 v2 and #50194-2024070410279309 v3).

For *in vivo* experiments, 6−12-week-old female mice were randomly assigned to specific treatment groups. All transgenic mice used were on a C57BL/6 background and were age-matched with wild-type controls for experiments.

### Animal procedures

MC38 mouse colon adenocarcinoma cell lines (kerafast, ENH204-FP) were cultured at 37 °C under 5% CO2 in the following culture medium: DMEM High Glucose containing 10 mM Hepes (Gibco, 11965092) supplemented with 10% (vol/vol) heat-inactivated (Hi) fetal bovine serum (Dutscher, 500105N1N, Batch S00G2), 1% Penicillin-Streptomycin-Amphotericin B Mix (PSA, PAN-Biotech, P06-07350), 1 mM Sodium Pyruvate (Gibco, 11360070), 2 mM L-glutamine (Gibco, 25030149) and MEM NEAA (0.1 mM each AA; Gibco, 11140050). Tumor cells (5×10^5^ MC38 cells per mouse) were resuspended in sterile Dulbecco’s Phosphate Buffer Saline (PBS, Gibco, 14190144) and implanted subcutaneously (s.c.) in mouse flank. Fourteen days after tumor engraftment, each mouse received one intratumoral (i.t.) injection of PBS containing 250 µg of DMXAA (referred to as DMX) or DMSO equivalent and tumors were harvested 5 h after treatment.

### *Ex vivo* procedures

#### Tumor infiltrating lymphocyte (TIL) analysis

MC38 tumors were subjected to mechanic dissociation and enzymatic digestion using the mouse tumor Dissociation Kit (Miltenyi Biotec, 130-096-730) and the gentleMACS™ Dissociator (Miltenyi Biotec, Germany) according to the manufacturer’s instructions. The single cell suspension was then washed two times using RPMI-1640 with L-Glutamine (Dutscher, 509090) supplemented with 10% Hi FBS, 1% PSA and 10 mM Hepes (Gibco, 15630080) - hereafter referred to as Complete RPMI - and passed through 70-µm and 30-µm cell strainers (Miltenyi Biotec, 130-110-916 and 130-110-915). Dead cells were then removed using the Dead Cell Removal Kit (Miltenyi Biotec, 130-090-101), and CD4 were enriched using mouse CD4 (TIL) MicroBeads, (Miltenyi Biotec, 130-116-475) according to the manufacturer’s instructions.

#### FACS analysis

Cells were then stained using Fixable Viability Dye eFluor™ 780 (eBioscience, 65-0865-18) for 20 min. Cells were then washed and stained with CD4-APC antibody (BD Bioscience, 553051) for 20 min. Finally, cells were stained for LC3 using Guava® Autophagy LC3 Antibody-based Detection Kit (Luminex, FCCH100171) following manufacturer’s instructions. Cells were analyzed using BD LSR II cytometer equipped with BD FACSDiva software (BD Biosciences, USA), and data were analyzed using FlowLogic software. Flow cytometry experiments were performed at the ImaFlow core facility part of the US58 BioSanD (Dijon, France).

### Cells

#### Primary BMDCs

Bone marrow cells were obtained from the femurs and tibias from mice. Red blood cells were lysed in Red Blood Cells lysis buffer (Invitrogen, 12770000) To generate BMDCs, the cells were cultured in complete RPMI supplemented with 50 µM of 2-Mercaptoethanol (Gibco, 21985023), 20 ng.mL^-1^ of GM-CSF (R&D systems, 415-ML-010/CF) and 10 ng.mL^-1^ of IL-4 (Miltenyi Biotec, 130-097-757) for 6 days with fresh medium supplementation at day 3. BMDCs were collected and plated in complete RPMI with 2-Mercaptoethanol before stimulation with STING agonists.

#### Primary CD4^+^ T cells magnetic enrichment

Total CD4^+^ T cells were enriched from mouse spleen and lymph nodes with mouse CD4 (L3T4) MicroBeads (Miltenyi Biotec, 130-117-043) according to the manufacturer’s instructions. Isolated CD4^+^ T cells were routinely 90% pure. Cells were seeded on plate-bound anti-CD3 (2 µg.mL^-1^; Clone 145-2C11; Bio X Cell, BE0001-1) and anti-CD28 (2 µg.mL^-1^; clone PV1; Bio X Cell, BE0015-5) antibodies in complete RPMI supplemented with 1 mM Sodium Pyruvate, 2 mM L-glutamine, MEM NEAA (0.1 mM each AA) and 50 µM 2-Mercaptoethanol.

#### Cell lines

WT and modified RAW264.7 mouse macrophages were cultured in DMEM supplemented with 10% Hi FBS and 1% PSA. WT and modified DC2.4 mouse dendritic cells (Merck, SCC142) were cultured in complete RPMI supplemented with 2 mM L-glutamine, MEM NEAA (0.1 mM each AA; Gibco), and 50 µM 2-Mercaptoethanol.

#### Primary Human PBMCs

Buffy coats of fresh blood from three healthy donors were obtained from the Etablissement Français du Sang (EFS) Bourgogne-Franche-Comté. In order to separate peripheral blood mononuclear cells (PBMC), buffy coats were diluted in PBS (1:1, v/v) and loaded on the top of Ficoll (Eurobio, CMSMCLO1). After 20 min of centrifugation at 700 × g (without deceleration), PBMCs were collected and washed twice with PBS and seeded in DMEM/F12 (Gibco, 31331093) without FBS. After 2 h, the cells were washed twice, and adherent myeloid cells were cultivated in DMEM/F12 supplemented with 10% FBS and 100 ng.mL^-1^ GM-CSF (Miltenyi Biotec, 130-093-866) for 6 days with medium change every 2 days.

### Genome editing by CRISPR Cas9

CRISPR-Cas9-based genome editing experiments were performed with the CRIGEN (CRISPR Functional Genomics Facility part of the US58 BioSanD). RAW 264.7 mouse macrophages and DC2.4 mouse dendritic cells were transfected with gRNA ribonucleoprotein (RNP) complex formed with sgRNA and Alt-R S.p. Cas9-GFP (Cas9-GFP:gRNA ratio used = 1 :1.2; Integrated DNA Technology/IDT, 1081058) with or without HDR donor oligo, using Lipofectamine^TM^ CRISPRMAX^TM^ Cas9 Transfection Reagent (Invitrogen^TM^, CMAX00001) according to the manufacturers’ instructions. The sequences of gRNA and donor oligos are provided in **Table S1**. One day after transfection, highly GFP-positive cells were FACS-sorted and single-cell seeded into each well of 96-well plates. After clonal expansion, knock-out (KO) clones were selected based on the absence of ATG13 detection by western blot and knock-in (KI) clones based on the detection of *Atg16l1* mutation by Sanger sequencing of PCR products.

### Cell Treatment

Cells were treated with 50 µg.mL^-1^ of 2’3’-cGAMP (cyclic [G (2’,5’)pA (3’,5’)p]) referred to as cG, Invivogen, tlrl-nacga23-1) or 5 µg.mL^-1^ of DMXAA (Murine STING ligand - Xanthenone Analog referred to as DMX, Invivogen, tlrl-dmx) for the indicated times in complete culture medium. Alternatively, cells were transfected with 50 µg.mL^-1^ of cdi-AMP (Bis- (3’-5’)-cyclic dimeric adenosine monophosphate, Invivogen, tlrl-nacda) or cdi-GMP (Bis-(3’-5’)-cyclic dimeric guanosine monophosphate, Invivogen, tlrl-nacdg) using Opti-MEM Glutamax (Gibco, 31985062) and Lipofectamine 2000 (Invitrogen, 11668019), according to the manufacturer’s instructions. TBK1/IKKε inhibition was performed by pre-treating cells overnight with 0.5 µM of BX795 (TBK1/IKKε inhibitor - InvitroFit™ Invivogen, tlr-bx7) before STING agonist addition for 3 h. To distinguish between autophagy and CASM, cells were pre-treated with Bafilomycin A1 (BafA1, 100 nM, Tocris, 1334) or PIK3C3-IN1 (1 µM, MedChemExpress, HY-12795) for 1 h and co-treated with cG for 3 additional hours. The inability of *Atg13* KO RAW264.7 and *ΔFip200* CD4 T^+^ cells to form autophagosomes was validated by treating cells with BafA1 (100 nM) or Monensin (Mon, 50 µM, Sigma, M5273) for 1 h or overnight with the mTOR inhibitor PP242 (5 µM, 4257; Tocris). The CASM-incompetence of DC2.4 ATG16L1^K490A^ cells was validated by treating cells PIK3CK (5 µM) for 2 h with or without BafA1 (100 nM) or Mon (50 µM) during the last hour.

### Western blotting

Total protein extracts were obtained by lysing cells into ice-cold RIPA Buffer (Pierce, 89900) containing Protease Inhibitor Cocktail (Sigma, P8340) and Phosphatase Inhibitor Cocktail 3 (Sigma, P0044) and Universal Nuclease (Pierce, 88700) on ice for 20 min. Lysates were centrifuged at 13,000 x g, 4°C for 10 min and supernatants were retrieved for protein quantitation (Pierce, 23225 or Bio-Rad, 5000111) followed by western blot analyses. Protein lysates were separated on 4-15% gradient precast gels (Bio-rad, 4561083) or home-made 10 or 15% gels before transfer on polyvinylidene difluoride (PVDF) membranes using a MIXED MW program on the Trans-Blot Turbo Transfer System (Bio-rad, 1704150) and kit (Bio-rad, 1704272) or by regular wet blotting system in Tris/Glycine buffer containing 20% of ethanol. Membranes were blocked in Tris-buffered saline (TBS, 20 mM Tris and 150 mM NaCl) containing 0.1% Tween 20 (Euromedex, 2001-B) (TBS-T) and supplemented with 5% bovine serum albumin (BSA, SEQUENS, 1005-70) or non-fat dry milk for 1 h at room temperature and incubated overnight at 4°C with primary antibodies (listed in **Table S2**) diluted in the appropriate blocking buffer. They were then incubated with anti-mouse or anti-rabbit horseradish peroxidase-conjugated secondary antibody (Cell Signaling Technology, 7076S and 7074S) diluted 1:5000 in TBS-T 5% BSA and proteins were detected with enhanced chemiluminescence (Clarity/Clarity Max Western ECL Substrate, Bio-rad, 1705060/1705062 or SuperSignal™ West Pico PLUS chemiluminescent, Pierce, 34580) using the ChemiDoc Imaging System (Bio-rad, 12003154) or the iBright CL750 Imaging System (Invitrogen, A44116). Densitometry analysis was performed using ImageJ software.

### RNA isolation/ RT-qPCR

Total RNA was extracted from cells with Quick RNA Microprep (Zymo Research, R1050) and retrotranscribed using iScript cDNA Synthesis Kit (Biorad, 1708891). cDNA were analyzed by real-time quantitative PCR (RT-qPCR) with PowerUp™ SYBR™ Green Master Mix (Applied Biosystems, A25778) according to the manufacturer’s instructions combined with 500 nM of Forward and Reverse primers (Primer sequences are indicated in **Table S1**) using the ViiA 7 Real-Time PCR System (Applied Biosystems)

### RNAseq

Total RNA was extracted from BMDCs using TRIzol™ Reagent (Invitrogen, 15596026) and subjected to DNAse treatment (Macherey-Nagel, 740963) and clean-up (Macherey-Nagel, 740948250) before determination of RNA quantity with QuantiFluor® RNA System (Promega, E3310) and quality with Bioanalyzer RNA 6000 Nano kit (Agilent, 5067-1511). Libraries were prepared from 300 ng of total RNA using PolyA mRNA retrotranscription with SuperScript IV Reverse Transcriptase (Invitrogen, 18090050) and TruSeq Stranded mRNA Library Prep (Illumina, 20020594), quantified using Quant-iT™ PicoGreen™ kit (Invitrogen, P7589) and checked for quality with Bioanalyzer High Sensitivity DNA Analysis kit (Agilent, 5067-4626) by ACTAgen facility (Université Paris-Saclay, France). Sequencing was done with single-end 75 bp reads using NextSeq 500/550 High Output Kit v2 and NextSeq NB552053 (Illumina, Texas, USA) by I2BC Next Generation Sequencing Platform (Gif-sur-Yvette, France). Raw FASTQ files were trimmed for residual adapter sequences, demultiplexed and quality filtered through bcl2fastq2-2.18.12, Cutadapt 3.2 and FastQC v0.11.5 respectively. Reads were pseudoaligned against GRCm38 - mm10 through Rsubread (Liao et al., 2019) package and differential expression analysis was performed using EdgeR (Chen et al., 2025). Gene set enrichment analysis was performed using fgsea R package (Korotkevich et al., 2021) and hallmark (H), ontology (C5), and curated (C2) gene sets from MSigDB mouse collections (https://www.gsea-msigdb.org/gsea/msigdb/collections.jsp). Specific pathways were selected. PPInfer R package (Jung & Ge, 2025) was used for representing GSEA barplots. For each of these pathways, signed difference ratios (SDR) of corresponding genes were calculated similar to Ramsey et al (Ramsey et al., 2008). Genes were then clustered based on these normalized profiles of expression. Clustering on rows was performed using Euclidian distance and complete agglomeration method. Statistical analyses were performed using the R software (http://www.R-project.org/), version 4.4.0, and graphs were drawn using GraphPad Prism V.9.0.2.

### ELISA & Cytokine Array

Cell culture supernatants were analyzed by ELISA for mouse IFN-β High Sensitivity ELISA Kit (pbl Assay Science, 42410) for T cell & BMDCs or DuoSet ELISA R&D (Systems/Bio-techne, DY8234-05) for RAW264.7 cells; mouse IL-6 and CXCL10 DuoSet ELISA (R&D Systems/Bio-techne, DY406-05 and DY466-05 respectively) or mouse TNF-α BD OptEIA™ ELISA (BD Bioscience, 560478). Cytokine Array analysis was performed using Proteome Profiler Mouse Cytokine Array Kit, Panel A (R&D Systems/Bio-techne, ARY006).

### Immunofluorescence

RAW 264.7 macrophages and DC2.4 cells_were grown on glass coverslips, incubated as described in the figure legends, fixed in ice-cold methanol at −20°C for 5 min and washed in phosphate-buffered saline (PBS; Sigma, D8537). Samples were blocked in 5% bovine serum albumin in PBS for 1 h at room temperature before overnight incubation with anti-LC3A/B (1:100; Cell Signaling Technology, 4108) in blocking buffer at 4°C. Following PBS washes, samples were incubated with Alexa Fluor 488 goat anti-rabbit (H+L) or Alexa Fluor 568 donkey anti-rabbit (H+L) secondary antibodies (1:500; Invitrogen, A-11034 and A-10042) for 1 h at RT. DNA was stained with DAPI (1 µg/mL; Sigma, D8417) before mounting coverslips with Fluoromount-G Mounting Medium (Invitrogen, 00-4958-02). Image acquisition was performed with the Axio Imager.M2 (Carl Zeiss Ltd, Germany) equipped with a 20X objective using Zen software Version 3.8 (Carl Zeiss Ltd). Quantification of LC3 dots per cell was performed with Icy software Version 2.5.3.0 (https://icy.bioimageanalysis.org). Microscopy experiments were performed at the ImaFlow core facility part of the US58 BioSanD (Dijon, France).

### Statistics

Statistical analyses were performed using Prism software (Graph Pad software, California, USA). For two-group comparisons, Unpaired t-tests was used. For multiple group comparisons, ordinary two-way ANOVA with Uncorrected Fisher’s LSD multiple comparisons test was used. P values < 0.05 were considered significant.

## Data availability statement

Raw RNAseq data used in this paper are available on SRA repository under BioProject accession number PRJNA1286784

## Acknowledgements

This work was possible thanks to the support of several facility staff. We thank Valérie Saint-Giorgio and all staff in the Plateforme de Zootechnie de l’Université Bourgogne Europe (Dijon, France). We thank E. Dubus, A. Bataille, N. Pernet and S. Monnier in the Imaflow facility (part of the US58 BioSanD, Dijon, France) for the help with data acquisition and technical advice. We thank Cyril Catelain in the Imaging and Cytometry facility (Gustave Roussy, Villejuif, France). This work has benefited from the facilities and expertise of the high throughput sequencing core facility of I2BC (Centre de Recherche de Gif – http://www.i2bc.paris-saclay.fr/). We thank Florent Dumont from the UMS-IPSIT BIOINFO facility who created the pipeline used for RNAseq data analysis, all staff in the ACTAGen core facility and the Région Ile-de-France for financial support to this facility. We would like to thank N. Ktistakis for critical feedback in preparation of this manuscript. We thank A. Baguet and M. Guittaut for providing DC2.4 cells and undergraduate students D. Genin, K. Kunene and V. Gomes Morais for their early work on STING agonists and *Atg13* KO cell lines respectively.

## Disclosure statement

Lionel Apetoh is a consultant for Brenus-Pharma. Lionel Apetoh performed consultancy work for Roche, Merck, Bristol-Myers Squibb, and Orega Biotech and was a recipient of a research grant from Sanofi. Oliver Florey reported non-financial support from Casma Therapeutics during the conduct of the study; personal fees from Casma Therapeutics outside the submitted work. No potential conflict of interest was reported by other authors regarding the submitted work.

## Funding

The authors were supported by grants from the Fondation pour la Recherche Médicale ARF20170938687 (EJ), the Conseil Régional de Bourgogne Franche-Comté and FEDER (LA, EJ and AR), the Agence Nationale de la Recherche (ANR-11-LABX-0021, LA and CP; ANR-24-CE17-1031, EJ; ANR-19-CE15-0028, PS-P and FG); The Fondation ARC DOC20190509200 (IB-L), the Embassy of France in the USA, 2018-2019 STEM Chateaubriand Fellowship (SHS), the Department of Defense Era of Hope Scholar Award BC200206/W81XWH-20-BCRP-EOHS (ERN), Keith W. and Sara M. Kelley Endowed Professor of Immunophysiology, University of Illinois Urbana Champaign (ERN), the Cancer Center at Illinois, University of Illinois Urbana Champaign (ERN) and the Biotechnology and Biological Sciences Research Council grants BB/Y006925/1 and BB/R019258/1 (OF, KH). This project has received funding from the European Research Council under the European Union’s Horizon 2020 research and innovation program (grant agreement number 677251), the Fondation de l’Avenir (AP-RM-22-020), the Agence Nationale de la Recherche (ANR-20-IDEES-0002) and the Ligue Nationale contre le cancer (CCIR Est and Comité des Hauts de Seine). This work was supported by the Institut National de la Santé et de la Recherche Médicale and in part by the Brown Center for Immunotherapy at Indiana University Melvin and Bren Simon Comprehensive Cancer Center.

## Abbreviations

ATG: autophagy related
ATG16L1: autophagy related 16 like 1
BMDCs: bone-marrow derived dendritic cells
CASM: conjugation of ATG8 to single membranes
2’3’-cGAMP/cG: cyclic guanosine monophosphate”adenosine monophosphate
CXCL10: C-X-C motif chemokine ligand 10
DMSO: dimethyl sulfoxide
DMXAA/DMX: 5,6-dimethylxanthenone-4-acetic acid
FIP200: FAK family kinase interacting protein of 200 kDa
IFN: interferons
IRF3: interferon regulatory factor 3
ISGs: IFN-stimulated genes
MAP1LC3B/LC3: microtubule associated protein 1 light chain 3
MFI: mean fluorescence intensity
NF-κB: nuclear factor kappa B
PBMC: peripheral blood mononuclear cell
PIK3C3/VPS34: phosphatidylinositol 3-kinase catalytic subunit type 3
RB1CC1: RB1-inducible coiled-coil protein 1
RTqPCR: real-time quantitative polymerase chain reaction
RUBICON: RUN and cysteine rich domain containing beclin 1 interacting protein
STING: stimulator of interferons genes
TBK1: TANK binding kinase 1
TILS: tumor infiltrating lymphocytes
TME: tumor microenvironment
TNF: tumor necrosis factor
WT: wild type

