## Supplemental Figures and Tables for "CASM potentiates STING-driven NFκB signaling in immune cells"

### Supplementary Files

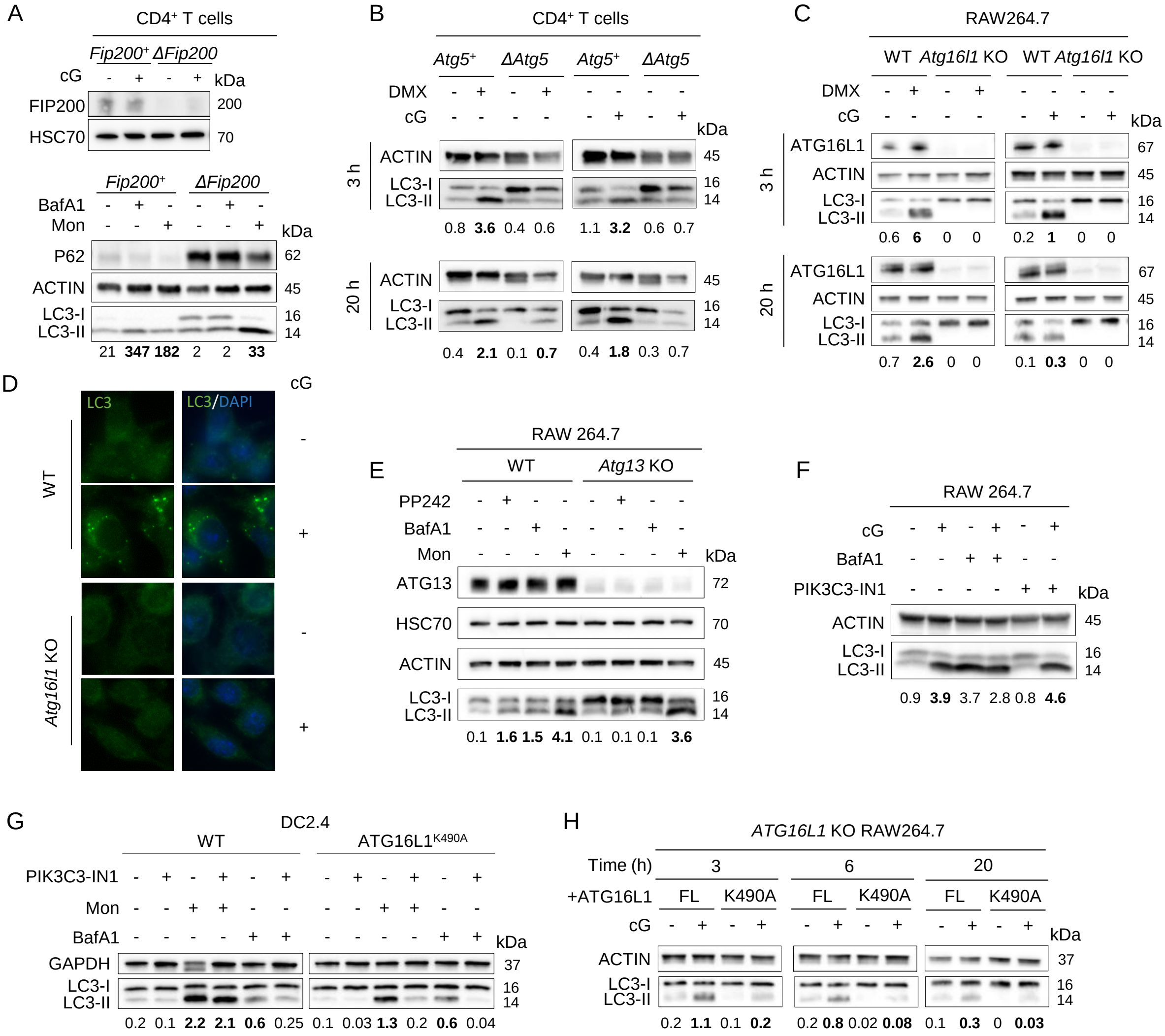

**Figure S1.** STING-induced LC3-lipidation requires components of LC3 lipidation machinery. (**A, upper panel**) FIP200 detection by western blot in *Fip200*⁺ or Δ*Fip200* CD4⁺ T stimulated or not with 2’3’-cGAMP (cG) for 3 h. LC3 lipidation analyzed by western blot (**A lower panel, B, C, E F, G and H**) or immunofluorescence (**D**) in *Fip200*⁺ or Δ*Fip200* CD4⁺ T (**A**) cells stimulated or not with Bafilomycin A1 (BafA1, 1 h) or Monensin (Mon, 1 h), representative of n= 3 independent experiments; in *Atg5*⁺ or Δ*Atg5* CD4⁺ T cells (**B**) stimulated or not with DMXAA (DMX) or cG for 3 or 20 h, representative of n=2 independent experiments; in WT or *Atg16l1* KO RAW264.7 macrophages (**C and D**), stimulated or not with DMX (**C**) or cG (**C and D**) for 3 (**C and D**) or 20 h (**C**), representative of n=3 (**C**) or n=2 (**D**) independent experiments; in WT or *Atg13* KO RAW264.7 macrophages (**E**), stimulated or not with Bafilomycin A1 (1 h), Monensin (1 h) or PP242 (overnight), representative of n=2 independent experiments; in RAW264.7 macrophages (**F**), stimulated or not with cG for 3 h with or without PI3KC3/VPS34 inhibitor (PI3KC3-IN1) or Bafilomycin A1 (1 h pretreatment and 3 h cotreatment) representative of n=3 experiments;  in WT or ATG16L1$^{K490A}$ DC2.4 cells (**G**), treated or not with PI3KC3-IN1, Monensin or Bafilomycin A1 for 1 h, representative of n=3 independent experiments; in Atg16l1 KO RAW264.7 macrophages (**H**) expressing exogenous Full-Length (FL) or K490A-mutated ATG16L1 (K490A), stimulated or not with cG for 3, 6 or 20 h, representative of n=3 independent experiments. LC3-II:LC3-I ratios are indicated below western blot images and increases in LC3-II:LC3-I ratios, compared to control conditions, are highlighted in bold.

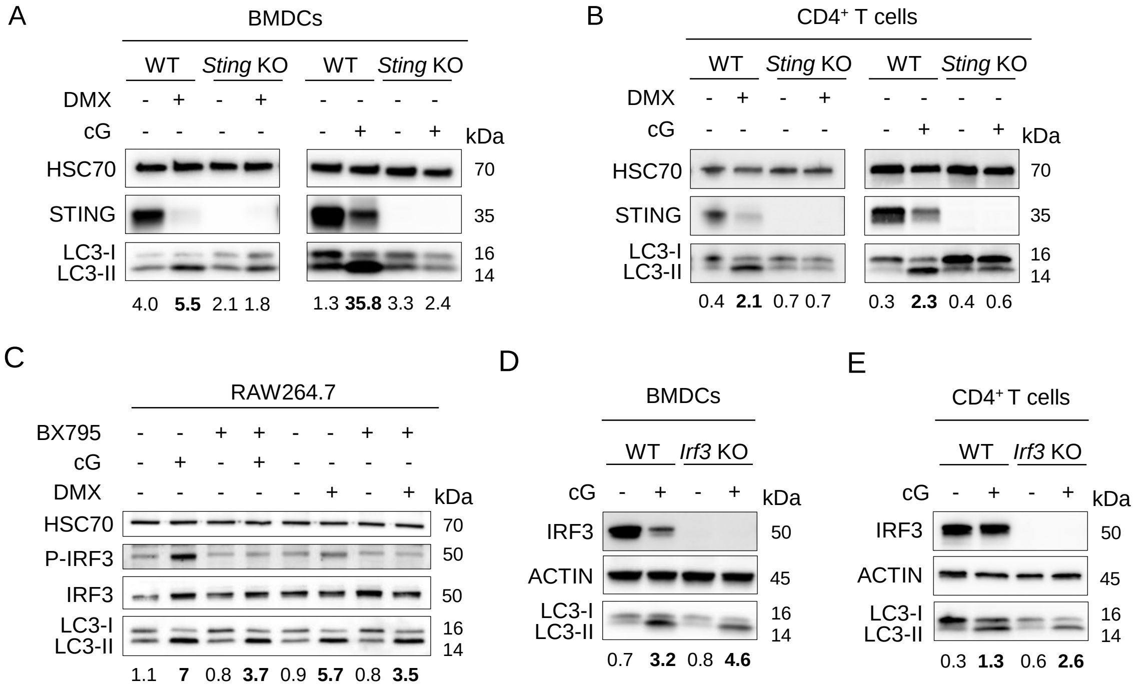


**Figure S2.** STING-driven CASM requires STING but not TBK1 and IRF3. LC3-II lipidation analyzed by western blot (**A to E**) in WT or *Sting* KO BMDCs (**A**) or CD4$^+$ T cells (**B**), stimulated or not with DMX or cG for 20 h, representative of n=2 or 3 independent experiments; in RAW264.7 macrophages (**C**) stimulated or not with cG or DMX for 3 h with or without overnight pretreatment with the TBK1 inhibitor BX795, representative of n=3 independent experiments;  in WT or *Irf3* KO BMDCs (**D**) or CD4$^+$ T cells (**E**), stimulated or not with cG for 20 h , representative of n=3 independent experiments. LC3-II:LC3-I ratios are indicated below western blot images and increases in LC3-II:LC3-I ratios, compared to control conditions, are highlighted in bold.

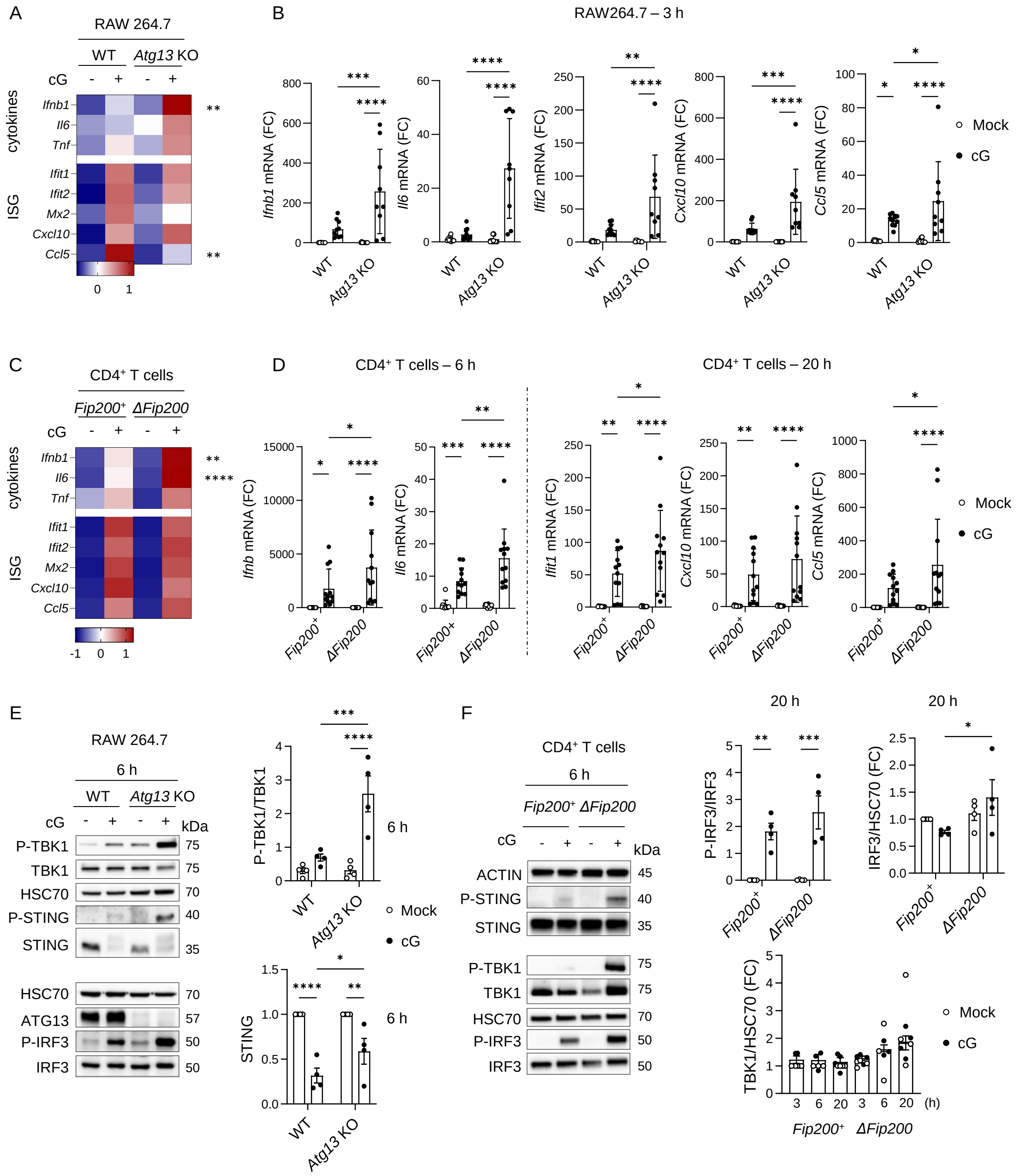


**Figure S3.** Autophagy differentially regulates the STING pathway in macrophages and T cells. Heatmap representing indicated cytokine and ISG mRNA expression (Z-score) determined by RTqPCR (A and C) from WT and *Atg13* KO RAW264.7 macrophages (**A**) or *Fip200*$^+$ and Δ*Fip200* CD4$^+$ T cells (**C**), stimulated or not with cG for 6 h, pooled data from 3 (**A**) or 4 (**C**) independent experiments, P values (**p<0.01, ****p<0.0001) determined by two-way ANOVA from each target relative expression; *Ifnb, Il6, Ifit1* (**B**) or Ifit2 (**D**), *Cxcl10* and *Ccl5* mRNA expression (Fold Change; Normalized to mean control condition from each genotype) determined by RTqPCR (**B and D**) from WT and *Atg13* KO RAW264.7 macrophages (**B**) or

*Fip200*$^+$ and $\Delta$*Fip200* CD4$^+$ T cells (**D**), stimulated or not with cG for 3 (**B**), 6 or 20 h (**D**), pooled replicates data from 4 independent experiments, P values (*p<0.05, **p<0.01, ***p<0.001, ****p<0.0001) determined by two-way ANOVA; western blot analysis of indicated proteins and corresponding relevant quantifications (**E and F**) from WT and *Atg13* KO RAW264.7 macrophages (**E**) or *Fip200*$^+$ and $\Delta$*Fip200* CD4$^+$ T cells (**F**) stimulated or not with cG for indicated time, pooled data (Mean +/- SEM) from 3 to 5 independent experiments, P values (*p<0.05, **p<0.01, ***p<0.001, ****p<0.0001) determined by two-way ANOVA.

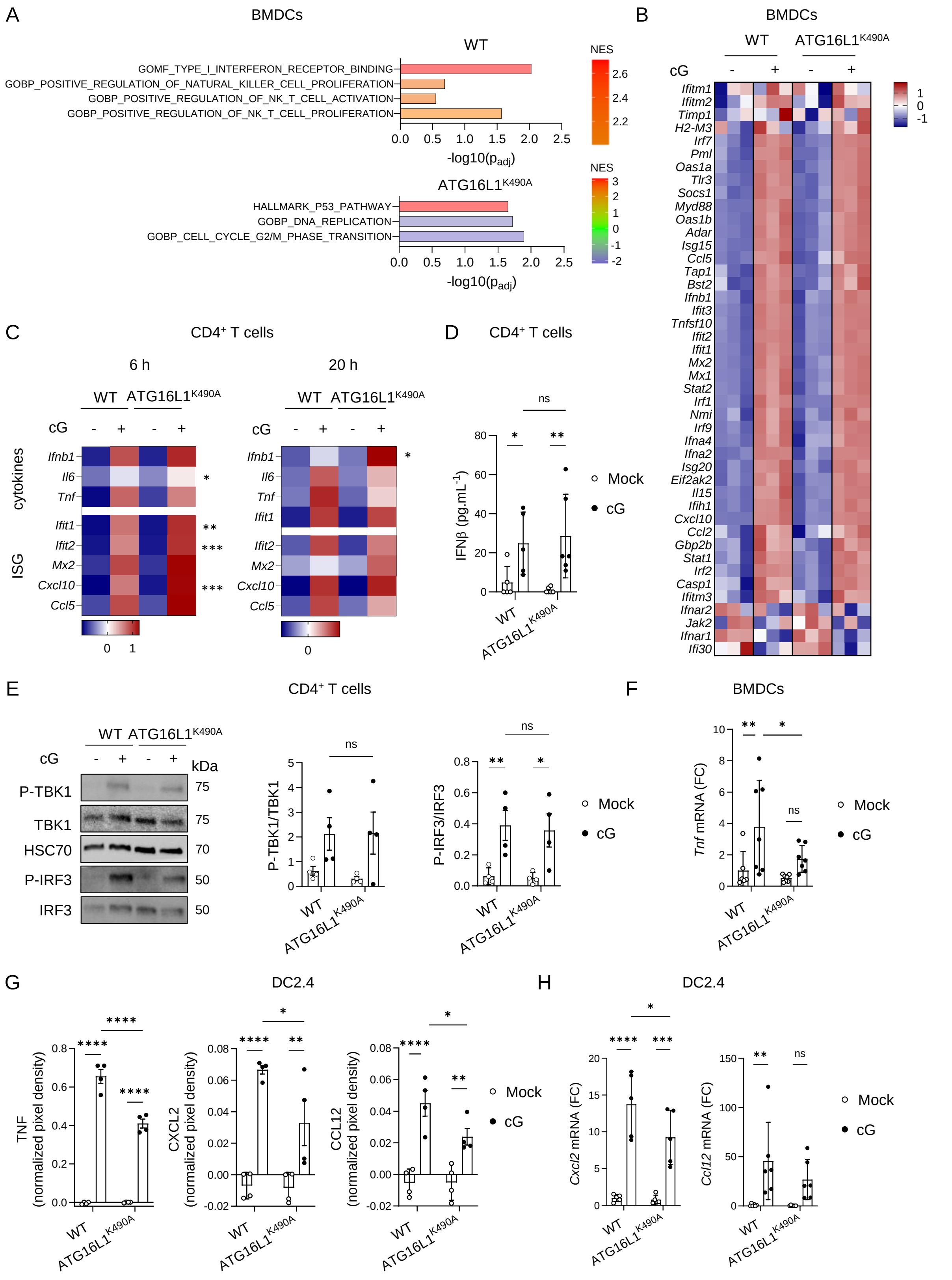


**Figure S4.** STING-driven CASM does not modulate the IRF3/IFN/ISG pathway but rather fine-tunes other inflammatory molecules in immune cells. (**A**) GSEA (Gene set enrichment analysis; Normalized Enrichment Score (NES) & -log10($p_{adj}$)) obtained from RNA sequencing analysis of WT or ATG16L1$^{K490A}$ BMDCs stimulated or not with cG for 20 h. (**B**) Heatmap representing RNAseq profiling of Type I IFNs (signed difference ratio (SDR) values) from WT or ATG16L1$^{K490A}$ BMDCs stimulated or not with cG for 3 h, columns represent samples, and each row represents one gene (**C**) Heatmap representing indicated cytokine and ISG mRNA expression (Z-score) determined by RTqPCR from WT or ATG16L1$^{K490A}$ CD4$^{+}$ T cells stimulated or not with cG for 6 or 20 h, pooled data (Mean) from n=3-4 mice from 2 independent experiments, P values (*p<0.05, **p<0.01, ***p<0.001) determined by two-way ANOVA from each target relative expression; (**D**) IFN-β secretion measured by ELISA from WT or ATG16L1$^{K490A}$ CD4$^{+}$ T cells stimulated or not with cG for 20 h**,** pooled data (Mean +/- SD) from n=5 mice from 3 independent experiments, P values (*p<0.05, **p<0.01) determined by two-way ANOVA. (**E**) Western blot analysis of indicated proteins and corresponding relevant quantifications, from WT or ATG16L1$^{K490A}$ CD4$^{+}$ T cells stimulated or not with cG for 3 h, pooled data (Mean +/- SEM) from n=4 mice from 3 independent experiments, P values (*p<0.05, **p<0.01) determined by two-way ANOVA. (**F**) *Tnf* mRNA expression (Fold Change FC; Normalized to mean of WT mouse control condition) determined by RTqPCR from WT or ATG16L1$^{K490A}$ BMDCs stimulated or not with cG for 3 h, pooled data (Mean +/- SD) from n=6-7 mice from 3 independent experiments, P values (*p<0.05, **p<0.01) determined by two-way ANOVA. (**G)** TNF, CXCL2 and CCL12 secretion measured by cytokine array from WT or ATG16L1$^{K490A}$ DC2.4 cells stimulated or not with cG for 6 h, pooled data (Mean +/- SEM) from 2 independent experiments, P values (*p<0.05, **p<0.01, ****p<0.0001) determined by two-way ANOVA. (**H)** *Cxcl2* and *Ccl12* mRNA (Fold Change FC; Normalized to mean of WT mouse control condition) from WT or ATG16L1$^{K490A}$ DC2.4 cells stimulated or not with cG for 6 or 3 h respectively**,** pooled data (Mean +/- SD) from 5 or 6 independent experiments, P values (*p<0.05, **p<0.01, ***p<0.001, ****p<0.0001) determined by two-way ANOVA.

### Supplementary Table

**Table S1.** Primer sequences

| | | |
|---|---|---|
| **Oligonucleotides used for CRISPR-based genome editing** | | |
| **Target/Type** | | **Sequence (5'-3')** |
| Mm *Atg13* crRNA | | caatcggttgtatacagtgtagg |
| Mm *Atg16l1* crRNA | | gttagggaagatcactgctc |
| Mm *Atg16l1* HDR donor | | taggtcagagagtgtggtccgagagatggaactgttagg ggcgatcactgccctggacctaaaccctgagagaactg agctcctgagctg |
| Mm *Atg16l1* Sequencing Forward primer (5'-3') | | ttcctgccccttgaagtctt |
| Mm *Atg16l1* Sequencing Reverse primer (5'-3') | | cctcagactcactcctgcaa |
| **Primers used for mouse gene expression quantification** | | |
| **Target** | **Forward Primer (5'-3')** | **Reverse Primer (5'-3')** |
| *Ifnb1* | ctccagctccaagaaaggac | tggcaaaggcagtgtaactc |
| *Il6* | ccagttgccttcttgggact | ggtctgttgggagtggtatcc |
| *Tnf* | agggtctgggccatagaact | ccaccacgctcttctgtctac |
| *Ifit1* | tcaaggcaggtttctgagga | attctctcccatggttgctgt |
| *Ifit2* | gctctggaaaaggacccgaa | gcttcagtgccaagaggact |
| *Mx2* | cctattcaccaggctccgaa | cagcataacctttttgcgaaattct |
| *Cxcl10* | cctatggccctcattctcac | ctcatcctgctgggtctgag |
| *Ccl5* | gtgcccacgtcaaggagtat | ttccttcgagtgacaaacacga |
| *Cxcl2* | tgaacaaaggcaaggctaactg | caggtacgatccaggcttcc |
| *Ccl12* | Bio-Rad, Primer Assay 10025636, ID qMmuCED0061017 | |
| *Gapdh* | aaggtcggtgtgaacggatt | attctcggccttgactgtgc |
| *Actin* | tgttaccaactgggacgaca | ctgggtcatcttttcacggt |

**Table S2.** Antibody list

| **Antibody (Clone)** | **Supplier** | **Reference** |
|---|---|---|
| anti-LC3A/B | Cell Signaling Technology | 4108 |
| anti-ATG13 (D4P1K) | Cell Signaling Technology | 13273 |
| anti-ATG16L1 (D6D5) | Cell Signaling Technology | 8089 |
| anti-TMEM173/STING | ProteinTech | 19851-1-AP |
| anti-Phospho-STING (Ser365) (D8F4W) | Cell Signaling Technology | 72971 |
| anti-HSC70 (B-6) | Santa Cruz Biotechnology | sc7298 |
| HRP-linked anti-ACTIN | Cell Signaling Technology | 12262 |
| anti-IRF3 (D83B9) | Cell Signaling Technology | 4302 |
| anti-Phospho IRF3 (Ser396) (4D4G) | Cell Signaling Technology | 4947 |
| anti-TBK1 (D1B4) | Cell Signaling Technology | 3504 |
| anti-Phospho TBK1/NAK (Ser172) (D52C2) | Cell Signaling Technology | 5483 |
| anti-NF-κB p65 (D14E12) | Cell Signaling Technology | 8242 |
| anti-Phospho-NF-κB p65 (Ser536) (93H1) | Cell Signaling Technology | 3033 |
| HRP-linked anti-GAPDH | Cell Signaling Technology | 3683 |
| anti-p62/SQTM1 | Progen | GP62-C |
| YY1 (D5D9Z) | Cell Signaling Technology | 46395 |
